# Fast and flexible Bayesian Jolly Seber models and application to populations with transients

**DOI:** 10.1101/2023.05.25.542303

**Authors:** James F. Saracco, Charles B. Yackulic

## Abstract

1. Jolly Seber (JS) models are an appealing class of capture-recapture models for modeling open populations because they allow for inferences about an array of population parameters, including abundance, survival, and recruitment. Multiple formulations of JS models have been developed and include both maximum likelihood and Bayesian approaches. Bayesian approaches offer greater flexibility; however, they are often extremely slow when applied to moderate to large populations because the latent states of all observed individuals, as well as any potential unobserved individuals (parameter-expanded data augmentation), must be simulated at each time step.
2. JS model implementations can be sped up dramatically through marginalization, whereby the conditional likelihoods of states are tracked for unique capture histories, rather than the simulated states of all individuals. Here we describe a marginalized implementation of a multistate JS model and compare its performance to that of an implementation using simulated discrete states. We fit models to data generated under two scenarios, one in which no data was missing and another in which 25% of data was randomly removed. To illustrate how marginalization can accommodate more complex models and datasets, we describe a modified version of the model for application to populations with transients and fit the model to simulated data and to data collected on Kentucky Warblers (*Geothlypis formosa*) as part of the Monitoring Avian Productivity and Survivorship (MAPS) program.
3. Both the discrete latent state and marginalized JS model implementations performed similarly with respect to bias and coverage; however, the marginalized implementation was roughly 1,000 times faster when no data were missing and 100 times faster when 25% of data was missing. The marginalized model accommodating transients also converged quickly and a more complex version of the model applied to the larger multi-site Kentucky Warbler data set yielded useful estimates of most parameters within a reasonable time frame (hours).
4. Gains in efficiency provided by marginalization shown here should stimulate additional study and development of this useful class of models and provide new opportunities for real-world applications to large complex data sets.

## Introduction

Open-population capture-recapture models are important tools for estimating demographic parameters of wildlife populations (Nichols 2005). These models include two basic classes: (1) Cormack-Jolly-Seber (CJS) models, which condition on first capture and provide estimates of apparent survival probability (Cormack 1964); and (2) Jolly-Seber (JS) models (Jolly 1965, Seber 1965), which are informed by entire capture histories (i.e., including occasions both prior to and after first capture). JS models are appealing because they make more complete use of available data and can provide inferences on an expanded set of demographic parameters including abundance and recruitment-related parameters. JS models assume that marked and unmarked animals have equal capture probability, which may limit their applicability in some situations (e.g., mark-resight studies). However, this assumption is likely reasonable for a broad array of studies that implement the same sampling protocol for both captures and recaptures (e.g., Robinson et al. 2009).

A variety of formulations of both CJS and JS models have been developed and include both maximum likelihood and Bayesian approaches (Pradel 1996, Schwarz and Arnason 1996, Link and Barker 2005, Dupuis and Schwarz 2007, Schofield and Barker 2008, Royle and Dorazio 2012, Tenan et al. 2014, Hostetter et al. 2021). Bayesian implementations offer several advantages over maximum likelihood methods, including ability to incorporate prior information, flexibility for handling hierarchical random effects, and relative ease with which derived quantities and their uncertainties can be calculated. A common Bayesian approach, known as state-space modeling or discrete latent state modeling (Royle 2008, Yackulic et al. 2020), involves simulating the state of individuals (e.g., alive/dead) and the conditional probability of their detection based on Bernoulli or categorical models applied to a set of observed capture histories. For JS models, the data set must be augmented to include a set of capture histories with no detections that represent individuals never observed to allow for modeling of both the survival and entry processes (Kéry and Schaub 2011). A drawback of this approach, however, is that computation times using Markov Chain Monte Carlo (MCMC) techniques can be prohibitive for many applications, particularly when the augmented data set includes many individuals or levels of stratification (e.g., sites or spatial strata).

Bayesian open-population models with discrete latent states can be sped up dramatically through marginalization (Yackulic et al. 2020). Rather than tracking the state of all individuals at each time step, these marginalized models only require tracking likelihoods of possible states given the demographic parameters for a reduced data set of unique capture histories and their frequencies. Yackulic et al. (2020) provide a thorough introduction to the marginalization in Bayesian population models, as well as code and examples for CJS models. Here we extend this work in the context of JS models. We focus on the multistate expression of the JS model described in Royle and Dorazio (2012), which is readily adaptable to a variety of model extensions involving additional states (Kéry and Schaub 2011). We apply the model to simulated complete data sets and data sets with random missing data, a common scenario in multi-site multi-collaborator data sets (e.g., Saracco et al. 2010), and compare parameter estimates and run times between the marginalized models and discrete latent state implementations using the popular open-source software program JAGS (Plummer 2003).

We also describe an extension of the basic marginalized multistate JS model to accommodate situations where not all sampled individuals are part of the resident populations. This model variant can both improve inferences about survival, recruitment, and population size, as well as allow estimation of residency probability and proportions of residents in populations (or, conversely, transience probability and proportions of transients), which may also often be of ecological interest (Wilson et al. 2018, Genovart and Pradel 2019, Oro and Doak 2020). Accounting for negative bias in survival estimates due to transiency, and estimation of residency or transiency probabilities, has been treated within the context of CJS models (Pradel et al. 1997, Nott and DeSante 2002, Hines et al. 2003, Royle and Dorazio 2008, Saracco et al. 2010). However, transiency or residency probability estimates from these models only apply to newly marked individuals and so may not be representative of the larger population because they depend, to some extent, on the proportions of previously marked individuals still alive and available for capture (Hines et al. 2003). Estimation of parameters related to the dynamics of resident and transient individuals within seasons has been developed within an ‘open robust design’ framework (Ruiz-Gutierrez et al. 2016); however, these models require additional data and model complication that may be unnecessary if primary interest is on between season dynamics. Treatment of these parameters in the JS context represents an important advance because it allows for disentangling of season specific observation (first capture) and state (residency) processes within a simpler multi season open population model.

We apply this ‘transient JS’ model to simulated data as well as to regional data from a migratory bird species, Kentucky Warbler (*Geothlypis formosa*), from bird-banding stations operated as part of the Monitoring Avian Productivity and Survivorship (MAPS) program (DeSante et al. 2004) in the Central Hardwoods Bird Conservation Region of the United States (Sauer et al. 2003).

## Materials and Methods

### Multistate Jolly Seber models: structure and data

Both the discrete latent state and multistate JS model assume a “superpopulation” (*sensu* Schwarz and Arnason 1996), *N*_super_, representing the number of individuals ever alive in the population. Membership in *N*_super_ is determined as a function of parameters describing the entry process and an augmented data set consisting of *M* individuals, including all those observed during the study, as well as any unobserved individuals that could have been members of the superpopulation. While the number of unobserved individuals to include in the augmented data set is somewhat arbitrary, the choice is not innocuous, particularly when using a discrete latent state implementation – too many and run times will be unnecessarily increased, too few and parameter estimates may be biased.

Following Kéry and Schaub (2011) and Royle and Dorazio (2012), we model the entry process in terms of a removal entry probability, γ, which represents the probability of entry for a member of *M* conditional on not having previously entered (see Appendix S1 for an alternative parameterization). Once entered, the probability of remaining alive and in the population is determined by a survival parameter, ϕ. We can represent the latent state of the *i* =1,…, *M* individuals across, t=0,…,*T* time periods with a matrix, *S*, with entries, *S*_i,t_ that can take values of: 1 = not yet entered into the population, 2 = alive and in the population, and 3 = dead. The number of time periods, *T*, includes all observed sampling occasions as well as a dummy occasion at *t =* 0 to allow modeling of initial entry into the population. All *S*_i,0_ are set equal to 1 to indicate that none have entered prior to the first observed time period at *t =* 1. The hidden Markov process determining *S*_i,t_ can be represented with an array, Ω, with individual entries ω_j,k,t_ representing probabilities of transitioning from latent state *j* (rows) in time *t* to latent state *k* (columns) in time *t*+1:

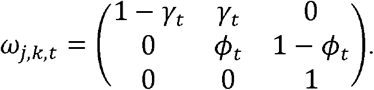

The data informing the latent states consist of a set of observed and unobserved (i.e., augmented) capture histories and, for the discrete latent state model, can be represented with a *M* × *T* matrix, **Y**, with entries *y*_*i,t*_. For the marginalized model, the *y*_*i,t*_ are summarized to a more compact matrix, **YS**, with entries *ys*_m,t_ that represent *m*=1,…,*MS* unique capture histories over the *T* years. The data informing the marginalized model are then the *ys*_m,t_ and their observed frequencies. The entries of **Y** and **YS** can take values of 1 (captured) or 2s (not captured). For computational purposes, we set all *y*_*i,O*_ =2and *ys*_*m,O*_ =2 for the initial dummy capture occasion. These capture histories are linked to latent states by a second array, **P**, with entries, *ρ*_j,o,t_ indexed by latent state *j* (rows), observed state *o* (columns), and period of observation *t* (not counting the dummy capture occasion), with time-specific capture probabilities represented by *p*_*t*_:

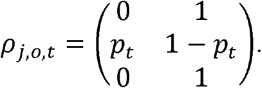

The principal difference between the discrete latent state and marginalized JS model implementations is the definition of the likelihoods. In the discrete latent state model, the true latent state is simulated for each individual and occasion for all *t* > 0based on a categorical model, *s*_i,t_ ∼Cat(∼_z[i,t-1],1:3,t-1_). The likelihood is then determined based on a categorical model of the observations conditional on the latent state, *y*_i,t_ ∼Cat(ρ_s[i,t],1:2,t_)

The likelihood for the marginalized JS model is defined in terms of an array, **Z**, which are defined (in order) by unique capture histories, *t*=0,…,*T* sampling occasions (again, tracks capture history likelihoods conditional on the demographic parameters. Its dimensions including the dummy occasion at *t*=0), and *s*-1:3states. Each entry, ζ_m,t,s_ is the likelihood that an individual with capture history *m* was in state *s* at time *t* conditional on the observed capture history through time *t* and values of the process and observation parameters.

As in the discrete latent state model, no individuals have entered the population at *t*=0. Thus, we set ζ_m,O,1_ =1and ζ_m,O,2:3_ =0 so that all individuals are available prior to the study.

For each subsequent time step, ζ_m,t,s_ is determined by first taking the inner product of ζ_m,t-1,1:3_ and the column vector taken from *s*^th^ column of the state transition matrix and then multiplying this quantity by the probability of the observed state given the state *s* (i.e., ζ_m,t,s_ = ⟨ ζ_m,t-1,1:3_, ω_1:3,s,t-1_ ⟩ × ρ _s,o[m,t],t_. For example, at the first true capture period (*t =* 1), the observation probability for capture histories with capture at that time period (i.e.,*y*_m,1_ =1) would be calculated as ζ_m,1,1_ =ζ_m,O,1_ ω_1,1,O_ ρ _1,1,1_ =1×(1-γ_O_)×0=0for individuals that did not enter (i.e., *s*=1); ζ_m,1,2_ =ζ_m,O,1_ω_1,2,O_ρ_2,1,1_ =1×γ_O_ ×*p*_1_ =γ_O_*p*_1_ for individuals that entered (i.e.,, =2); and ζ_m,1,3_ =ζ_m,O,1_ζ_1,3,O_ρ_2,1,1_ =1×0×0=0for dead individuals (i.e., *s* = 3). Thus, the only possible transition for individuals captured at the first true capture occasion is entry into the population.

The likelihood of each capture history is determined by summing the likelihoods of all possible states, *s*, after the final capture occasion, 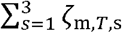, weighted by the frequency of the *m*^th^ capture history. In BUGS language, as implemented in popular software programs such as WinBUGS (Lunn et al. 2000), OpenBUGS (Lunn et al. 2009), JAGS (Plummer 2003), and NIMBLE (de Valpine et al. 2017), a modified version of the “ones trick” can be used to update the posterior with these values. See Yackulic et al. (2020) and Appendix S1 for detail; Yackulic et al. (2020) also provide the appropriate approach to updating the posterior using the program Stan (Carpenter et al. 2017).

For both the discrete latent state and marginalized models, unconditional time-specific the entry probabilities from the superpopulation to the population, *b*_t_ are derived as a function of entry probabilities conditional on not having previously entered, *c*_t_, where *c*_O_ =γ_O_ and 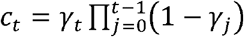for and the inclusion probability (i.e., probability of being a member of the superpopulation), ψ (Kéry and Schaub 2011): *b*_t_ =*c*_t_/ ψ. In discrete latent state models, secondary parameters (i.e., derived quantities) can be calculated directly from the modeled latent states. For example, time-specific numbers of newly entered individuals (i.e., newborns or immigrants), *B*_t_, can be derived by counting the Z_i,t_ =2|Z_i,t-1_ =1, and total time-specific abundance, *N*_t_, can be calculated by counting the Z_i,t_ =2for each sample of the posterior distribution.

Alternatively, secondary parameters can be derived as functions of primary (i.e., modeled) parameters, other secondary parameters, and/or the input data; and this approach can be applied to either model type. For example, numbers of new entrants can be estimated as *B*_t_ =*C*_t_*M*, and the superpopulation size, *N*_super_, can then be derived as 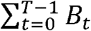 (Schwarz and Arnason 1996). Time-specific abundance can be estimated as *N*_1_ =*B*_O_ for the initial sampling period and *N*_t_ =*B*_t-1+_ϕ_t_*N*_t-1_ for *t*>1, or alternatively could be estimated with the canonical estimator,*N*_t_ =*n*_t_ /*p*_t_, where the *n*_t_ represents a count of the number of individuals observed alive at time *t* (i.e., summing the frequencies of all *y*_i,t_ =1or *ys*_m,t_ = 1). Other quantities of interest that can be derived include population growth rate, λ_t_ =*N*_t+1_/*N*_t_, and per capita entry rate, *f*_*t*_ =*B*_*t*_/*N*_*t*_ (or, alternatively, *f*_*t*_ =λ_*t*_ ϕ_t_; Pradel 1996, Link and Barker 2005). Additional methods for deriving secondary parameters from marginalized models are also possible. For example, annual abundance can be derived by simulating latent states from reconstituted individual capture histories (i.e., converting **YS** back to **Y**) and summing them in a manner analogous to what is described above for the discrete latent state models (see Appendix S2 for detail).

### Marginalized multistate Jolly Seber model with transients

Capture-recapture studies often include individuals that are sampled but are not resident in the local population. These transient individuals have been defined in the context of CJS models as individuals having zero probability of being alive and available for capture following initial capture (Pradel et al. 1997). In the JS context, however, transients can be defined more generally as individuals appearing only once and never returning to the sampling area. Thus, interpretation of transience and residency probabilities in the JS context is more ecologically relevant because it refers to the larger population, not just the population of newly marked individuals. We can accommodate transiency in the multistate JS model by adding a state that allows for entry as either a resident or a transient, such that there are now four possible states, including: 1 = not yet entered; 2 = alive resident; 3 = alive transient; and 4 = dead. Distinguishing between the two alive states requires an extra parameter π, denoting the probability of being a resident. Thus, individual entries in the transition and capture probability arrays become:

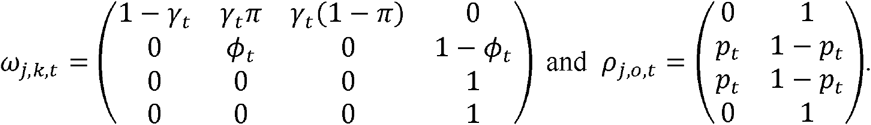

The residency probability parameter, π, is estimated in the marginalized multistate transient JS model based on 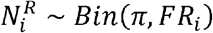, where 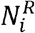is the number of residents represented by unique capture history *i*. In some cases, the residency state can be partially observed. For example, within a ‘robust design’ sampling framework (Pollock et al. 1990), whereby multiple secondary sampling occasions are completed within each primary sampling occasion, individuals observed on multiple secondary sampling occasions during the primary sampling occasion in which they were marked can be used as an indicator of residency (Nott and DeSante 2002, Hines et al. 2003). To accommodate this scenario, and following logic developed for the discrete latent state version of a transient CJS model (Saracco et al. 2010), we can define an observation model for residency as: 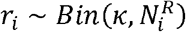, where *r*_i_ is the number of observed residents for capture history *i, k* is the probability of observing residency, and 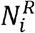 is the true number of residents for capture history *i*. Note that in contrast to the CJS scenario, where residency observation and state probabilities are specific to years of first capture and allow for temporal modeling, here both *K* and π are time-constant.

Nevertheless, we can derive numbers of transients and residents entering the population at each time step as 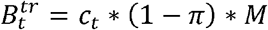 and 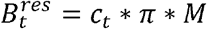 respectively. For transients, the number of entries is equivalent to the number of transients in the population at any time step; 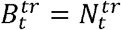. At *t*=1, the total number of residents is also the same as the number of resident entries, 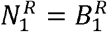; the number of residents can be derived as 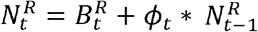 for subsequent time steps. The proportion of residents at any given time can then be derived as 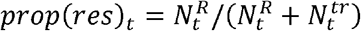.

A critical assumption of this formulation of the transient JS model is that *p*_t_, the probability of capture at least once across the season, is the same for both transient and resident individuals. This is plausible if most individuals enter the population early in the season and we think of *p*_t_ as an initial capture probability, π as the probability of persisting for multiple sampling periods, and κas the probability of capturing an individual that persists for multiple periods. Nevertheless, we acknowledge that in many cases residents and transients will have differential exposure to capture, resulting in violations of the assumption of equal *p*_t_ between these two groups. Accounting for these differences can be accommodated with additional model development and data from secondary samples within seasons (e.g., Ruiz-Gutierrez et al. 2016). For simplicity, we do not consider this additional complication here.

### Discrete v. marginalized multistate JS models: a simulation study

To compare the relative speed and efficacy of discrete latent state and marginalized JS models, we simulated 100 capture-recapture data sets. As in Kéry and Schaub (2011), we assumed a superpopulation size of *N*_super_ =400across *T* – 1 = 7 capture occasions, a constant survival probability of ϕ=0.70, entry probabilities of *b*_1_ =0.34and *b*_2. 7_ =0.11, and a constant capture probability of *p*=0.50. We assumed *U* (0,1) prior distributions for survival, ϕ, capture probability *p*, and entry probabilities *b*_t_. Data sets were augmented with 500 all-ones capture histories to account for unobserved individuals for a total of *M* capture histories (observed + unobserved). These individual capture histories made up the basic data sets for the discrete latent state JS model, an *M*×*T*,matrix, where *T* represents the number of sampling occasions, including the dummy occasion. For the marginalized model, data sets (including the augmented individuals) were aggregated to *MS* unique observed capture histories and their frequencies.

To explore potential effects of missing data, we randomly assigned 25% of the individual × capture occasion data generated in the simulated data sets to be missing. For the marginalized models, capture histories were then collapsed to unique capture history × effort (missing or not). Missing data in the two model types are handled differently. For the discrete latent state models, missing data values are imputed with the categorical models of the state and observation process in the same way they are for the complete data situations, which should result in relatively little increases in model run times. However, for the marginalized models, where state and capture probabilities are tracked with deterministic nodes, missing data can be accommodated by setting capture probabilities to zero for missed occasions. This requires tracking capture probabilities and conditional state likelihoods across an additional dimension allowing for different capture probabilities among individuals sampled at different times, which may result in relatively greater loss in efficiency. An alternative approach, which we do not explore here but that could also have implications for model run times, would be to define a multistate model for *p*_t_, whereby *p*_t_ can take ‘available’ or ‘unavailable’ states depending on whether sampling occurred (e.g., Lovich et al. 2014).

For the discrete latent state models, we derived secondary parameters as functions of the summed latent states as in Kéry and Schaub (2011). For the marginalized models, we derived secondary parameters as analytical functions of the primary parameters and data or other second-order states, as described above, as a simulation study based on the primary parameter values used here showed better credible interval coverage, faster implementation time, and less biased parameter estimates compared to alternative methods of deriving second-order states and secondary parameters (Appendix S2).

To assess models and data sets, we compared the ability of models to recover data generating values, 95% credible interval coverage, credible interval widths, and model run times. We implemented all models in JAGS (Plummer 2003) with the jagsUI package (Kellner 2015) in R (R Core Team 2021). R code for simulating data and implementing models is provided in the data package (Saracco and Yackulic 2023), and a detailed description of model code is provided in Appendix S1.

### Transient marginalized multistate JS models

We applied a transient marginalized multistate JS model to 100 simulated data sets using the same parameters as the marginalized JS model above, but with the added residency parameters set to π=0.6and *k*=0.4. Simulation and model details are provided in the data package.

We applied a version of this model to capture-recapture data from 2912 adult Kentucky Warblers (*Geothlypis formosa*) banded between 1993 and 2015 in the Central Hardwoods Bird Conservation Region in the United States (Sauer et al. 2003) as part of the Monitoring Avian Productivity and Survivorship (MAPS) program. These data were collected at 51 MAPS stations, which on average operated ∼ 10 yrs (sd = 6) of the 23-yr time span, resulting in ∼ 42% (28058 of 66976) of the individual × year data points missing. For the analysis, these data were summarized to yield 876 unique capture histories. We used a *Beta*(1/(*T*-1)2-*t*/(*T*-1))prior distribution on γ_t_ due to the missing data and large number of sampling periods (Dorazio 2020). Initial attempts using *U* (0,1)priors for γ_t_ with MAPS data resulted in marginal posterior distributions on N peaking at M, regardless of how many capture probability, we modeled *p*_sta[i]_ with a logit-linear model with intercept and a random unobserved individuals the data set was augmented with. To allow for spatial variation in station effect distributed as *Norm*(0, σ^2^). In addition, we included annual variation in survival by modeling ϕ_t_ with a logit-linear model with intercept and a random year effect distributed as *Norm*(0, σ^2^). We applied standard vague priors for intercepts (*Norm*(0,0.01) and variances of random effects (*Unif* (0,10).

## Results

### Discrete v. marginalized multistate JS models

Median estimates of the data generating parameter values were successfully recovered for both the discrete latent state and marginalized multistate JS model implementations (Fig. 1). Credible interval coverage was close to nominal at the 95% level for most parameters and scenarios, although it was slightly lower for the *N*_super_ and, parameters (Table 1; Fig. 1). Coverage for the *N*_super_ and, parameters for models with missing data was especially low (∼ 80-90%). The abundance estimates,*N*_t_ and *N*_super_, tended to be biased high for all models and data scenarios. Median bias of posterior medians for the abundance parameters was lowest for the discrete latent state model with complete data (+ 3-7%), slightly higher for the marginalized model with complete data (+ 5-8%), and greater still for the models with missing data (+ 6-12% for the discrete latent state model and + 7-14% for the marginalized model). Estimates of capture probability, *p*, tended to be biased low and was similar between models (median – 5% bias in posterior *p* medians for both models with complete data and – 6% for models with missing data). Credible interval widths were essentially identical between the discrete and marginalized models for the entry, survival, and capture probability parameters; however, widths were ∼ 10-20% greater for derived parameters, such as *N*_super_ (Table 1)The marginalized models ran much faster than the discrete latent state models, reaching 1000 effective samples for the least converged parameter ∼ 100 “(missing data model) to 1000 “faster than the discrete latent state versions (Fig. 2).

**Table 1.** 95% credible interval coverage and widths [mean with (SE) below] for the discrete latent state (D), marginalized (M), discrete latent te with 25% missing data (D.NA), and marginalized with 25% missing data (M.NA) multistate Jolly Seber models applied to 100 simulated a sets.

|  | % coverage |  |  |  |  | Width |  |  |  |
| --- | --- | --- | --- | --- | --- | --- | --- | --- | --- |
|  | D | M | D.NA | M.NA |  | D | M | D.NA | M.NA |
| $N_{super}$ | 90 (3.0) | 92 (2.7) | 83 (3.8) | 87 (3.4) | | 82.1 (0.9) | 99.3 (0.7) | 116.4 (1.7) | 129 (1.3) |
| $\phi$ | 94 (2.4) | 94 (2.4) | 92 (2.7) | 91 (2.9) | | 0.104 (0.000) | 0.104 (0.001) | 0.132 (0.001) | 0.132 (0.001) |
| $b_1$ | 98 (1.4) | 98 (1.4) | 97 (1.7) | 98 (1.4) | | 0.157 (0.001) | 0.157 (0.001) | 0.185 (0.001) | 0.185 (0.001) |
| $b_2$ | 97 (1.7) | 97 (1.7) | 97 (1.7) | 97 (1.7) | | 0.010 (0.010) | 0.010 (0.010) | 0.010 (0.010) | 0.010 (0.010) |
| $b_3$ | 96 (2.0) | 96 (2.0) | 98 (1.4) | 97 (1.7) | | 0.129 (0.001) | 0.129 (0.001) | 0.154 (0.002) | 0.155 (0.002) |
| $b_4$ | 94 (2.4) | 94 (2.4) | 96 (2.0) | 97 (1.7) | | 0.123 (0.001) | 0.124 (0.001) | 0.150 (0.002) | 0.150 (0.002) |
| $b_5$ | 95 (2.2) | 95 (2.2) | 96 (2.0) | 96 (2.0) | | 0.122 (0.001) | 0.122 (0.001) | 0.148 (0.002) | 0.148 (0.002) |
| $b_6$ | 95 (2.2) | 95 (2.2) | 97 (1.7) | 97 (1.7) | | 0.119 (0.001) | 0.119 (0.001) | 0.146 (0.002) | 0.146 (0.002) |
| $b_7$ | 95 (2.2) | 95 (2.2) | 96 (2.0) | 96 (2.0) | | 0.115 (0.001) | 0.115 (0.001) | 0.142 (0.002) | 0.142 (0.002) |
| $p$ | 93 (2.6) | 92 (2.7) | 87 (3.4) | 89 (3.1) | | 0.139 (0.001) | 0.140 (0.001) | 0.183 (0.002) | 0.184 (0.002) |

**Figure 1.**
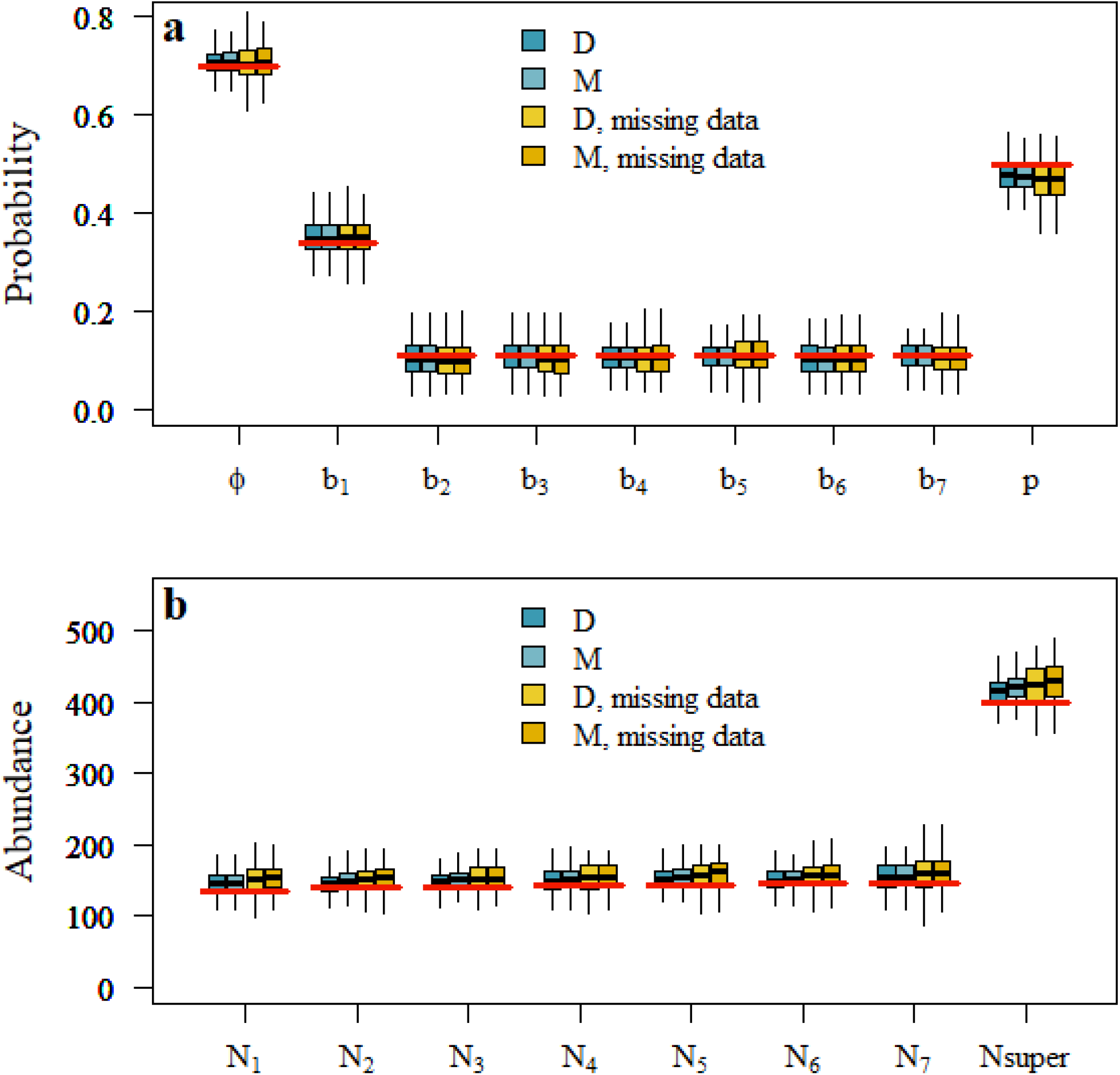
Boxplots of posterior medians for adult apparent survival probability, cp, entry probability, b, capture probability, *p* (a); annual abundance, N_t_, and superpopulation abundance, N_super_, (b) from discrete (D) and marginalized (M) multistate Jolly Seber models applied to 100 complete simulated data sets and 100 data sets with 25% missing data. Data generating values are indicated with red lines.

**Figure 2.**
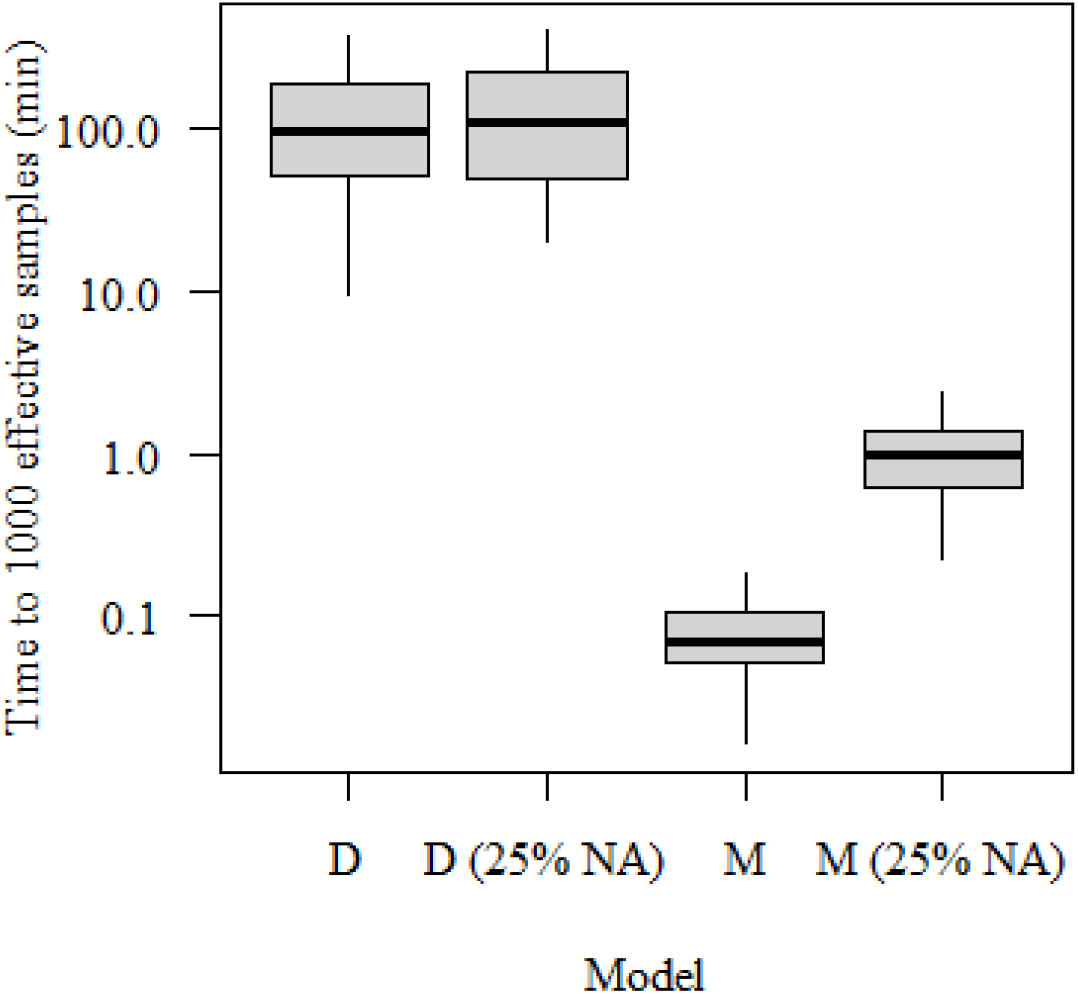
Boxplot of the time required to reach 1000 effective samples of the least converged parameter for discrete (D) and marginalized (M) multistate Jolly Seber models applied to 100 complete simulated data sets and to the same 100 data sets but with 25% of the data randomly missing.

### Transient model applied to simulated data

The model converged quickly, with a median time to achieve 1000 effective samples of the least converged parameter of 0.5 minutes. Parameter estimates were very similar to the data generating values, suggesting that the model can provide reliable inferences about residency probability, numbers of residents in the population, survival and recapture probabilities under the assumption of equal *p*_t_ between residents and transients. As for the basic model version, however, estimates of capture probability tended to be biased low and population size high (Fig. 3). Residency probability also tended to be biased high (Fig. 3a). Coverage of 95% credible intervals was relatively low for capture probability and superpopulation size of residents (between 80% and 85%, Fig. 3c).

**Figure 3.**
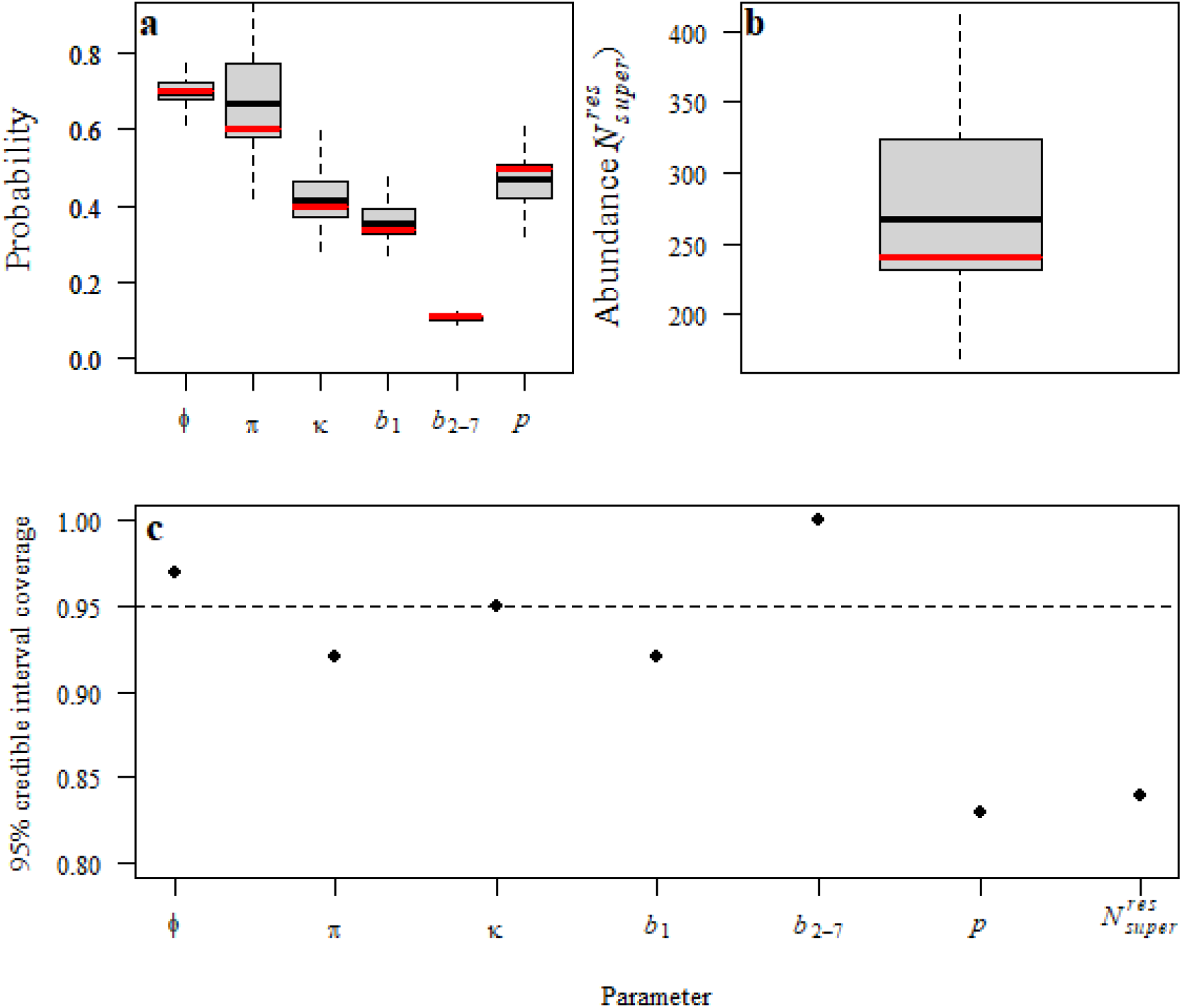
Boxplots of posterior medians for adult apparent survival probability, residency probability, entry probability, capture probability (a), and superpopulation size of resident individuals (b) from a marginalized multistate transient Jolly Seber model applied to 100 simulated data sets. Data-generating values are indicated with red lines. Mean 95% credible interval coverage of model parameters is shown in (c).

### Application to MAPS data

Based on a model run with 3 MCMC chains of 40000, burn-in of 10000, and thinning rate of 3 (for a total of 30000 saved posterior samples), the model took approximately 24 hours to run, considerably longer than our simpler simulation examples. Most parameters converged relatively quickly with the median time to achieve 1000 effective samples of 4 hours; however, a few parameters took longer with the least converged parameter taking 3.7 days to achieve 1000 effective samples. All parameters had 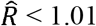, suggesting satisfactory convergence.

Model results suggested that survival probability was relatively consistent across the study (ϕ =0.5 [0.46, 0.54]; σ=0.09 [0, 0.3]; Fig. 4a). The derived recruitment rate was much more variable, particularly for the first and last time intervals (Fig. 4b). However, precision of the recruitment estimates was also lower for those intervals and likely reflect the low number of stations operated during the first and last years of the time series (8 and 3, respectively). Annual population change and abundance matched patterns of recruitment (Figs. 4c, d), suggesting the importance of this demographic rate in driving population dynamics. Residency probability, π was estimated to be (π = 0.53 [0.44, 0.64], and the proportion of residents in the population was relatively consistent across years, again with the exception of first and last years (Fig. 4e).

**Figure 4.**
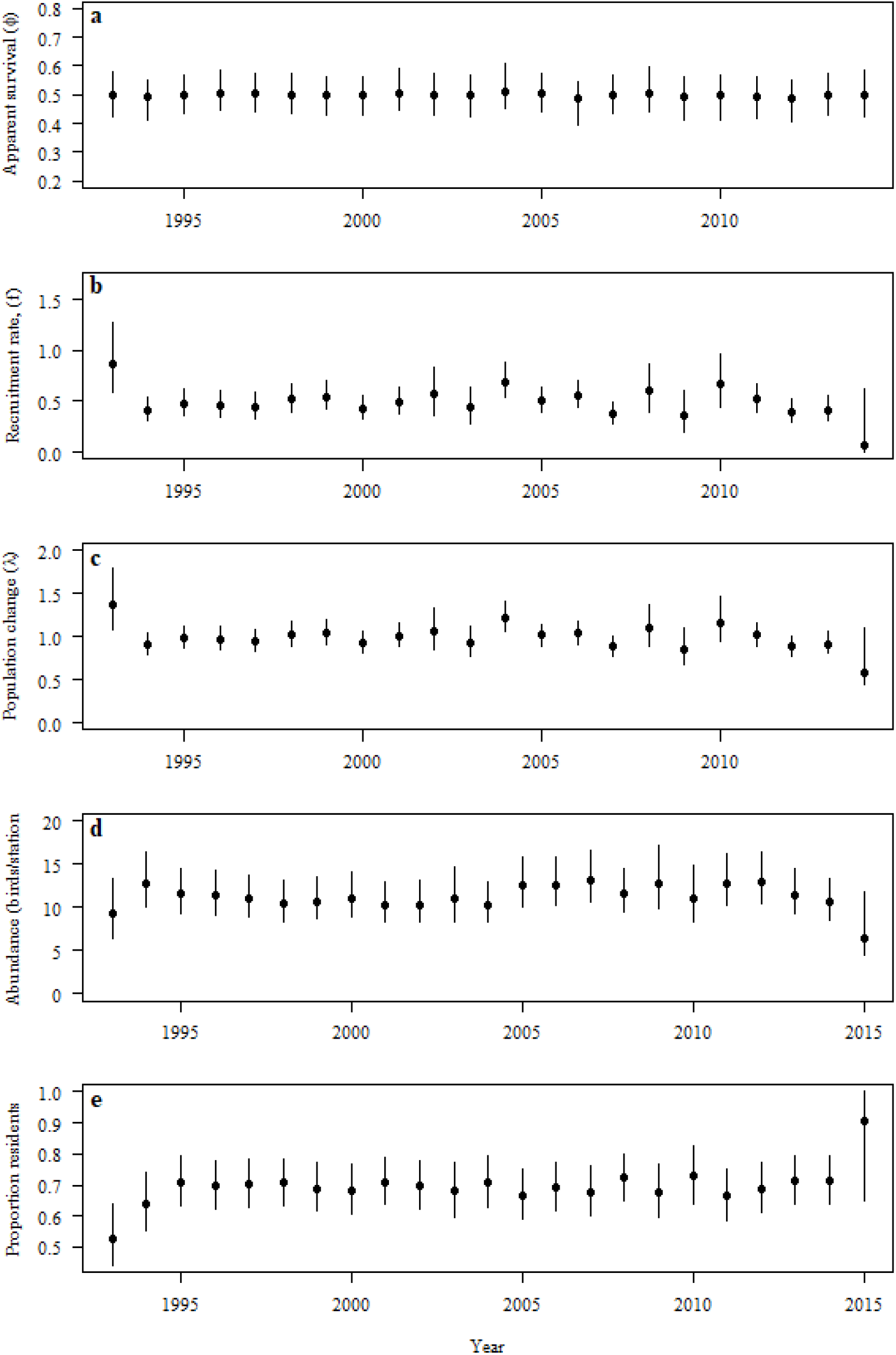
Medians + 95% credible intervals for annual demographic parameters estimated for Kentucky warblers banded in the Central Hardwoods Bird Conservation Region as part of the Monitoring Avian Productivity and Survivorship (MAPS) program.

## Discussion

Bayesian modeling with MCMC estimation provides a flexible framework for implementing capture-recapture models with latent states and complex hierarchical model structures. However, because common Bayesian implementations of these models require simulating discrete latent states of each individual at each time step, they can be slow to converge and require long run times for moderate to large data sets (Yackulic et al. 2020). This is especially true for JS models, where observed capture histories must be augmented with potentially large numbers of unobserved individuals to encompass the population of interest. Here we show that through marginalization, whereby the individual capture histories are aggregated to unique capture histories (or unique capture history × covariate) and their frequencies, and state probabilities, rather than individual states, are tracked, parameter estimates can be recovered in a fraction of the time. This gain in efficiency greatly expands the potential applications for this useful class of capture-recapture model.

Because JS models do not condition on first capture, they require an assumption of equal capture probability for captured and uncaptured individuals, which may limit their utility for certain sampling scenarios (e.g., mark-resighting studies). However, whenever this assumption is reasonable, these models provide an attractive alternative to CJS models, because they enable inferences about an expanded suite of demographic parameters including recruitment, population size, and population dynamics. As proposed in the original formulations of Jolly (1965) and Seber (1965), population size can be estimated within a CJS modeling context by simply dividing the number of individuals captured by the recapture probability estimate (Dugger et al. 2004, McDonald et al. 2005 p. 241, see Yackulic et al. 2014, Appendix C for a complex application). However, this approach can lead to population size estimates that suggest negative recruitment because entry parameters are not included in the likelihood (Schwarz and Arnason 1996). By modeling both the recruitment and survival processes, JS approaches provide a means of preventing such inconsistencies. Other options that do condition on capture, yet are still able to provide inferences on recruitment through modeling capture histories both forward and backward in time, offer an alternative approach (Pradel 1996, Link and Barker 2005). These ‘reverse-symmetry’ models offer an advantage over JS models in that the recruitment or population growth parameters can be included in the likelihood and so modeled directly, rather than derived from the survival and population size parameters (Tenan et al. 2014, Saracco et al. 2020). In addition, because they are based on applying time-specific multinomial likelihoods to a compact set of sufficient statistics, they too can be implemented quickly using Bayesian methods. Nevertheless, these reverse symmetry models do not allow the same level of flexibility as the multi-state JS models for handling complexities such as transiency or demographic rates that depend on states that change over time [e.g., age class or breeding status; Kendall et al. (2019), Hostetter et al. (2021)]. Finally, while these models can provide inference about population dynamics parameters, they are largely unable to inform population size (but see Tenan et al. 2019).

Median parameter estimates from our simulation study were very similar between the discrete latent state and marginalized JS models for the set of parameter values we considered in our simulations. However, both model implementations showed slight negative bias in capture probability estimates and positive bias in population size estimates. These biases were magnified for model runs with random missing data. We suggest that additional study is needed to better understand the potential effects of missing data, including different amounts and types of missing data (e.g., random v. consecutive missed years at particular sites), on inferences across a range of combinations of parameter values. Understanding the consequences of missing data is especially important for monitoring programs, such as MAPS, where as much as half the sampling occasions can commonly be missed due to stations entering and leaving the program over the years.

Credible intervals were wider for the marginalized v. discrete latent state models for the population size and other parameters derived as deterministic functions of population size. Nevertheless, these estimates exhibited better coverage compared to the discrete latent state model or marginalized model with parameters derived from conditional estimates applied to reconstituted capture histories (Appendix S2). Given the greater speed of implementation, better coverage, and lack of bias in year-specific population size, recruitment, and population change estimates, we suggest that this approach appears to be the most generally useful among those considered here for implementing models and estimating derived parameters.

We described a variation of the basic marginalized multistate JS model to illustrate the versatility of the basic modeling framework to address a common complication of many capture-mark-recapture programs that involves the sampling of individuals that are not part of the target population. This ‘transient’ version of the marginalized multistate JS model assumes equal capture probability between residents and transients, which may be unrealistic in many situations; however, because capture probability is informed here by between-year recaptures, we suggest that it is unlikely to bias estimates of resident population size. Higher resident than transient recapture probabilities would nonetheless be expected to result in underestimates of transient populations and overestimates of the proportion of residents in populations. Additional within-season data and model development (e.g., Ruiz Gutierrez et al. 2016) will be required to provide a more complete understanding of resident and transient components of populations.

The version of the transient JS model that we describe here successfully recovered parameter values from our simulations, although the same biases in capture probability and superpopulation estimates seen in the basic JS version were also evident in the transient model, and there was some evidence in bias in the residency probability parameter as well. We suggest that additional study is needed to assess how biases may change across different combinations of parameter values.

While our application to the MAPS Kentucky Warbler data set was much slower than the simple simulation scenarios we considered, this was not entirely unexpected given the larger dimensions of the data set (∼ 20 x the number of rows representing unique site x capture histories and > 3 x the number of sampling occasions compared to the simulated transient model data sets), random effects structure for the survival and capture probability models, and large number of missed sampling occasions. Even with the longer run time, reasonable inferences for most parameters were obtained within a few hours, demonstrating the potential utility of this modeling framework for more complicated real-world examples.

Ideally, study design, objectives, and characteristics of the data set at hand should dictate possible model applications without having to consider constraints of long implementation times. We hope that the marginalized versions of the JS model described here will stimulate additional study and development of this useful class of models and open new potential applications to large complex data sets.

## Acknowledgements

Funding was provided by the Knobloch Family Foundation. R. Siegel and S. Albert provided logistical support. We are grateful to the many MAPS cooperators and to D. DeSante for his work in developing and managing the MAPS program. D. Kaschube helped to prepare the MAPS data set with support from E. Cox, L. Helton, and R. Taylor. R. B. Taylor and W. Kendall provided helpful comments that improved the manuscript. Any use of trade, product, or firm names in this publication is for descriptive purposes only and does not imply endorsement by the U.S. Government. This is Contribution Number 731 of The Institute for Bird Populations.

## Author Contributions

J. Saracco and C. Yackulic conceived the ideas and designed methodology; J. Saracco prepared and analysed the data and led the writing of the manuscript. Both authors contributed critically to the drafts and gave final approval for publication.

## Data Availability

R code and data needed to reproduce results are archived in the Open Science Framework https://doi.org/10.17605/OSF.IO/Q3458 (Saracco and Yackulic 2023).

