## Appendix S1 for "Fast and flexible Bayesian Jolly Seber models and application to populations with transients"

**Appendix S1**. BUGS/JAGS code for multistate Jolly Seber models.

In our simulation studies, we compared the discrete multistate JS model based on code in Kéry and Schaub (2012), Ch. 10, and compared results to a marginalized version of the same model. Here we provide model code in BUGS language, as implemented in popular software programs such as OpenBUGS (Lunn et al. 2009), JAGS (Plummer 2003), or NIMBLE (de Valpine et al. 2017) for these two model variations. Additional model detail is provided in the main manuscript.

First, priors for survival ($\phi$), removal entry ($\gamma$), and capture probabilities ($p$) for both the discrete and marginalized models can be written as:

```
  for (t in 1:(n.occasions-1)){
    phi[t] <- mean.phi
    gamma[t] ~ dunif(0, 1)
    p[t] <- mean.p
  }

  mean.phi ~ dunif(0, 1)
  mean.p ~ dunif(0, 1)
```

Note that we follow Kéry and Schaub (2012) and Royle and Dorazio (2012) in parameterizing the recruitment process in terms of the removal entry probabilities, which can be related to the true entry probabilities, $b_{t}$, following:

$$b_{1}=\gamma_{1}/\psi\text{ for }t=1\text{ and }b_{t}=\gamma_{t}\prod_{j=1}^{t-1} \left( 1-\gamma_{j} \right)/\psi\text{ for }t>1,$$

where $\psi$ represents the inclusion probability (i.e., probability that a member of the augmented data set is part of the superpopulation). In BUGS language, this can be written as:

```
for (t in 1:(n.occasions-1)){
    qgamma[t] <- 1-gamma[t]
    b[t] <- c[t]/psi
  }
  for (t in 2:(n.occasions-1)){
    c[t] <- gamma[t]*prod(qgamma[1:(t-1)])
  }
psi <- sum(c[1:(n.occasions-1)])
c[1] <- gamma[1]
```

Alternatively, we could define a model for $b_{t}$ and derive the removal entry probabilities. One way of doing this would be modeling the $b_{t}$ with a Dirichlet prior distribution:

```
  b[1:(n.occasions-1)] ~ ddirich(alpha[1:(n.occasions-1)])
  psi ~ dunif (0, 1)
  gamma[1]<-psi*b[1]
 for (t in 2:(n.occasions-1)){
    gamma[t] <- psi*b[t]/(1-psi*sum(b[1:(t-1)]))
    }
```

Under this parameterization, the prior for the $\alpha_{t}$ parameters of the Dirichlet distribution must be entered as data for JAGS to run. Setting the $\alpha_{t}$ values to 1 yields an uninformative prior distribution with equal probabilities of entry at each time period. Limited initial testing suggests that this parameterization runs even faster in JAGS than does the parameterization with a model for $\gamma_{t}$ (approximately 60% faster with parameter values used in the paper and no missing data) while yielding identical results. The $b_{t}$ could also be modeled under a generalized linear modeling framework with a multinomial logit regression. For example, as:

for (t in 1:(n.occasions-2)){

b0[t] ~ dunif(0, 1)

lb0[t] <- log(b0[t]/(1-b0[t]))

logit(b[t]) <- lb0[t]

} # t

b[(n.occasions-1)] <- 1 - sum(b[1:(n.occasions-2)])

The prior for $\psi$ and equation for $\gamma_{t}$ remain the same as defined for the Dirichlet model above. Although this approach runs slower than the other parameterizations, it provides a means of modeling the entry probabilities as functions of covariates. Note that initial values for the $b_{t}$ should be set to relatively small values to avoid the ‘invalid parent values’ error in JAGS.

Second, state transition ($\boldsymbol{\Omega}$) and observation ($\mathbf{P}$) arrays for both the discrete latent state and marginalized model types can be represented with:

```
#-----------------------------
# States
# 1 = not yet entered
# 2 = alive
# 3 = dead

# Observations
# 1 = captured
# 2 = not captured
#-----------------------------

# omega tracks probability of state at t+1 given state at t
  for (t in 1:(n.occasions-1)){
    omega[1,1,t] <- 1-gamma[t]
    omega[1,2,t] <- gamma[t]
    omega[1,3,t] <- 0
    omega[2,1,t] <- 0
    omega[2,2,t] <- phi[t]
    omega[2,3,t] <- 1-phi[t]
    omega[3,1,t] <- 0
    omega[3,2,t] <- 0
    omega[3,3,t] <- 1

# rho tracks capture probability at t given state at t
    rho[1,1,t] <- 0
    rho[1,2,t] <- 1
    rho[2,1,t] <- p[t]
    rho[2,2,t] <- 1 - p[t]
    rho[3,1,t] <- 0
    rho[3,2,t] <- 1
  }
```

Note that Kéry and Schaub (2012) also tracked state and observation probabilities across a fourth dimension of $\boldsymbol{\Omega}$ and $\mathbf{P}$ representing individuals. This is unnecessary here, as parameters do not vary among individuals, and run time in JAGS (Plummer 2003) is approximately halved by not indexing by individuals. For the discrete latent state model, we then simulate the state ($s_{i,t}$) and conditional observations ($y_{i,t}$) from categorical distributions for all $i=1,\ldots,M$ individuals at each time step:

```
  for (i in 1:M){
    s[i,1] ~ dcat(1)
    for (t in 2:n.occasions){
      s[i,t] ~ dcat(omega[s[i,t-1],1:3,t-1])
      y[i,t] ~ dcat(rho[s[i,t],1:2,t-1])
    }
  }
```

Note that the indexing for rho[,,t-1] here refers to capture probabilities at time t.

For the marginalized model, we loop through the $i=1,\ldots,MS$ unique capture histories to track probabilities of state transitions and observations. Contributions of each unique capture history likelihood to the overall likelihood were determined in the BUGS model using the modified “ones trick” described by Yackulic et al. (2020), whereby capture history frequencies are provided as data for both the frequency response variable $\text{fr}$ and the number of trials $\text{FR}$ in binomial distributions with success probabilities defined by the likelihoods:

```
  for (i in 1:MS){
    zeta[i,1,1] <- 1  # all individuals unrecruited at first dummy occasion
    zeta[i,1,2] <- 0
    zeta[i,1,3] <- 0

    for (t in 2:n.occasions){
      zeta[i,t,1] <- inprod(zeta[i,(t-1),1:3], omega[1:3,1,(t-1)] * rho[1,ys[i,t],(t-1)])
      zeta[i,t,2] <- inprod(zeta[i,(t-1),1:3], omega[1:3,2,(t-1)] * rho[2,ys[i,t],(t-1)])
      zeta[i,t,3] <- inprod(zeta[i,(t-1),1:3], omega[1:3,3,(t-1)] * rho[3,ys[i,t],(t-1)])
    }
    lik[i] <- sum(zeta[i,n.occasions,1:3])
    fr[i] ~ dbin(lik[i], FR[i])
  }
```

For the discrete latent state model we can then derive secondary parameters from first-order states following:

```
for (i in 1:M){
   for (t in 2:n.occasions){
      al[i,t-1] <- equals(s[i,t],2)
      } # t
   for (t in 1:(n.occasions-1)){
      d[i,t] <- equals(s[i,t]-al[i,t],0)
      } #t
   alive[i] <- sum(al[i,])
   } # i

for (t in 1:(n.occasions-1)){
   N[t] <- sum(al[,t])        # population size
   B[t] <- sum(d[,t])         # number of entries. Note that indexing in paper
                              # goes from 0 (initial entry) to T (n.occasions - 1)
   } # t
for (i in 1:M){
   w[i] <- 1-equals(alive[i],0) # w[i] = 1 if individual i ever entered the
                                # population and 0 otherwise
   } # i
for(t in 1:(n.occasions-2)){
  lambda[t] <- N[t+1]/N[t] # population growth rate
  f[t] <- B[t+1]/N[t] # recruitment (per capita entry) rate. Could also be
                      # calculated as lambda[t] - phi[t]
} # t
Nsuper <- sum(w[]) # could also be calculated with sum(B[])
```

For the marginalized models, we can derive secondary parameters as functions of model parameters and first-order states:

```
  for (t in 1:(n.occasions-1)){
    qgamma[t] <- 1-gamma[t]
  } # t
  c[1] <- gamma[1]
  N[1] <- B[1]
  for (t in 2:(n.occasions-1)){
    c[t] <- gamma[t] * prod(qgamma[1:(t-1)])
    N[t] <- B[t] + phi[t] * N[t-1]
  } # t
  psi <- sum(c[1:(n.occasions-1)])    # inclusion probability
  for (t in 1:(n.occasions-1)){
    b[t] <- c[t] / psi      # entry probability
    B[t] <- c[t]*M          # number of entries
  } # t

  for(t in 1:(n.occasions-2)){
  lambda[t] <- N[t+1]/N[t] # population growth rate
  f[t] <- B[t+1]/N[t]      # recruitment (per capita entry) rate
  } # t

  Nsuper <- sum(B[1:(n.occasions-1)]) # superpopulation size
```

### Missing data models

While the code for the discrete latent state model does not change with missing data (which are just entered as “NA” values), the marginalized models require the following modifications. First, for capture probability we add an index for unique capture history and multiply by an indicator variable, $op.ind_{i,t}$ denoting whether the individuals represented by that capture history were observable ($op.ind_{i,t}=1$) or not ($op.ind_{i,t}=0$) during that time period:

for (t in 1:(n.occasions-1)){

$$\vdots$$

for (i in 1:MS){
 p[i,t] <- mean.p*op.ind[i,t] # op.ind: 1 = effort, 0 = no effort
 }

$$\vdots$$

}

Second, for $\mathbf{P}$ we need to add a fourth dimension for the capture probability array:

for (t in 1:(n.occasions-1)){

$$\vdots$$

```
    for (i in 1:MS){
    rho[1,1,t,i] <- 0
    rho[1,2,t,i] <- 1
    rho[2,1,t,i] <- p[i,t]
    rho[2,2,t,i] <- 1 - p[i,t]
    rho[3,1,t,i] <- 0
    rho[3,2,t,i] <- 1
    }
  }
```
