## Appendix S2 for "Fast and flexible Bayesian Jolly Seber models and application to populations with transients"

Supporting information for: Saracco, J. and C. B. Yackulic. Efficient and flexible Bayesian Jolly Seber models and application to populations with transients

**Appendix S2**. Variations of multistate Jolly Seber models and methods for estimating derived quantities

We investigated four methods for estimating derived secondary parameters but reported only the first two approaches in the manuscript. These methods include:

1) Estimating parameters using model “d”, the discrete latent state model presented in the manuscript. In this approach derived parameters are calculated directly from the posteriors of first-order states.
2) Estimating parameters using model “m1”, the marginalized model presented in the manuscript. In this approach derived parameters are calculated as functions of modeled parameters (see main manuscript and Appendix S1 for details).
3) Estimating parameters using model “m2”, which was not presented in the manuscript, but is identical to m1 except for in how derived parameters are estimated. In this approach, $N_{super}$ is calculated by estimating the number of unobserved individuals by drawing from a binomial distribution using the probability of inclusion and the number of augmented individuals and this estimate of unobserved individuals is added to the known number of observed individuals. Other derived parameters are calculated by determining the full conditional estimates of different underlying dynamics based on the observed capture history and simulating feasible true recruit and death timing based on their relative likelihood based on an observed capture history.
4) Estimating parameters using model “m3,” which was not presented in the manuscript, but is identical to m1 except for in how derived parameters are estimated. In this approach, $N_{super}$ is calculated using the same approach as “m2”. $N_{t}$ is derived by calculating forward-conditional probabilities for each individual and time period and using a series of Bernoulli trials to determine the expected number of alive individuals. B’s are calculated as functions of modelled parameters as in “m1”.

We compared these four methods with respect to speed, coverage, and precision of estimates of derived parameters based on 100 simulated data sets. Below we share annotated code, as well as figures based on these simulations.

Credible interval coverage for the year-specific abundance estimates, $N_{t}$ was relatively low for the discrete latent state model and especially the marginalized model m2 (Table S1). Credible interval coverage was highest and generally above the nominal 95% level for model m1, although this likely reflects the broader credible intervals for that model. For model m2, $N_{t}$ estimates were biased high for time periods 3-7 (Fig. S1). Model m1 was fastest, but all of the marginalized models were relatively fast compared to model d (Fig. S2).

***Table S1****. 95% credible interval coverage and widths [mean (sd)] for discrete (d) and marginalized (m1, m2, and m3) JS models applied to 100 simulated data sets that differed with respect to how abundance (N), population change, and recruitment are estimated.*

|  | d | | m1 | | m2 | | m3 | |
| --- | --- | --- | --- | --- | --- | --- | --- | --- |
|  | % coverage | Width | % coverage | Width | % coverage | Width | % coverage | Width |
| $N_{super}$ | 90.0 | 80.4 | 93.0 | 97.0 | 89.0 | 80.7 | 90.0 | 77.2 |
|  | (30.2) | (11.4) | (25.6) | (8.5) | (31.4) | (10.6) | (30.2) | (9.7) |
| $N_{1}$ | 65.0 | 68.4 | 72.0 | 81.5 | 85.0 | 55.1 | 63.0 | 74.6 |
|  | (47.9) | (9.1) | (45.1) | (8.6) | (35.9) | (6.4) | (48.5) | (11.4) |
| $N_{2}$ | 72.0 | 54.7 | 85.0 | 67.3 | 60.0 | 50.7 | 77.0 | 57.5 |
|  | (45.1) | (7.4) | (35.9) | (6.8) | (49.2) | (5.8) | (42.3) | (9.0) |
| $N_{3}$ | 79.0 | 51.2 | 90.0 | 63.7 | 54.0 | 51.5 | 85.0 | 57.1 |
|  | (40.9) | (7.0) | (30.2) | (6.5) | (50.1) | (5.6) | (35.9) | (8.2) |
| $N_{4}$ | 85.0 | 50.0 | 93.0 | 63.6 | 48.0 | 53.6 | 88.0 | 57.7 |
|  | (35.9) | (6.6) | (25.6) | (6.1) | (50.2) | (5.7) | (32.7) | (7.8) |
| $N_{5}$ | 85.0 | 50.2 | 95.0 | 64.0 | 52.0 | 55.3 | 88.0 | 57.7 |
|  | (35.9) | (7.2) | (21.9) | (6.5) | (50.2) | (6.6) | (32.7) | (9.1) |
| $N_{6}$ | 91.0 | 52.0 | 98.0 | 65.4 | 65.0 | 57.2 | 91.0 | 57.7 |
|  | (28.8) | (7.1) | (14.1) | (6.6) | (47.9) | (7.0) | (28.8) | (8.6) |
| $N_{7}$ | 90.0 | 56.1 | 93.0 | 67.2 | 86.0 | 64.4 | 89.0 | 56.6 |
|  | (30.2) | (7.9) | (25.6) | (7.6) | (34.9) | (9.6) | (31.4) | (8.3) |
| $\lambda_{1}$ | 84.0 | 0.403 | 89.0 | 0.451 | 87.0 | 0.406 | 79.0 | 0.429 |
|  | (36.8) | (0.078) | (31.4) | (0.098) | (33.8) | (0.072) | (40.9) | (0.087) |
| $\lambda_{2}$ | 90.0 | 0.367 | 92.0 | 0.410 | 88.0 | 0.314 | 84.0 | 0.412 |
|  | (30.2) | (0.057) | (27.3) | (0.071) | (32.7) | (0.042) | (36.8) | (0.063) |
| $\lambda_{3}$ | 93.0 | 0.353 | 95.0 | 0.392 | 90.0 | 0.281 | 93.0 | 0.401 |
|  | (25.6) | (0.043) | (21.9) | (0.051) | (30.2) | (0.029) | (25.6) | (0.047) |
| $\lambda_{4}$ | 87.0 | 0.339 | 91.0 | 0.374 | 87.0 | 0.260 | 90.0 | 0.386 |
|  | (33.8) | (0.050) | (28.8) | (0.060) | (33.8) | (0.029) | (30.2) | (0.055) |
| $\lambda_{5}$ | 88.0 | 0.343 | 90.0 | 0.376 | 80.0 | 0.253 | 88.0 | 0.390 |
|  | (32.7) | (0.051) | (30.2) | (0.062) | (40.2) | (0.026) | (32.7) | (0.058) |
| $\lambda_{6}$ | 89.0 | 0.354 | 92.0 | 0.385 | 85.0 | 0.268 | 87.0 | 0.390 |
|  | (31.4) | (0.048) | (27.3) | (0.061) | (35.9) | (0.025) | (33.8) | (0.052) |
| $f_{1}$ | 85.0 | 0.416 | 91.0 | 0.458 | 90.0 | 0.445 | 78.0 | 0.445 |
|  | (35.9) | (0.086) | (28.8) | (0.103) | (30.2) | (0.080) | (41.6) | (0.137) |
| $f_{2}$ | 90.0 | 0.369 | 93.0 | 0.414 | 89.0 | 0.342 | 88.0 | 0.433 |
|  | (30.2) | (0.059) | (25.6) | (0.074) | (31.4) | (0.045) | (32.7) | (0.083) |
| $f_{3}$ | 93.0 | 0.349 | 97.0 | 0.397 | 89.0 | 0.307 | 92.0 | 0.418 |
|  | (25.6) | (0.044) | (17.1) | (0.053) | (31.4) | (0.032) | (27.3) | (0.058) |
| $f_{4}$ | 88.0 | 0.332 | 92.0 | 0.379 | 85.0 | 0.290 | 89.0 | 0.393 |
|  | (32.7) | (0.052) | (27.3) | (0.065) | (35.9) | (0.036) | (31.4) | (0.074) |
| $f_{5}$ | 87.0 | 0.333 | 92.0 | 0.383 | 85.0 | 0.302 | 88.0 | 0.393 |
|  | (33.8) | (0.053) | (27.3) | (0.066) | (35.9) | (0.038) | (32.7) | (0.073) |
| $f_{6}$ | 90.0 | 0.342 | 92.0 | 0.393 | 86.0 | 0.326 | 89.0 | 0.392 |
|  | (30.2) | (0.051) | (27.3) | (0.064) | (34.9) | (0.034) | (31.4) | (0.063) |

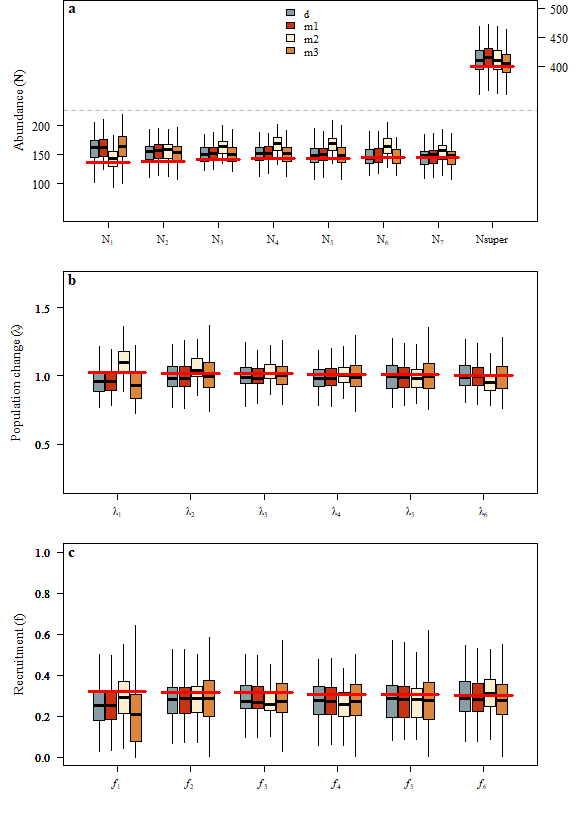

***Figure S1****. Boxplots of posterior medians of derived parameters from discrete (d) and marginalized (m1, m2, m3) multistate Jolly Seber models applied to 100 simulated data sets. Abundance estimates are shown in (a; note right y-axis shows scale for N_super_); population change in (b) and recruitment in (c). Data-generating values are indicated with red lines.*

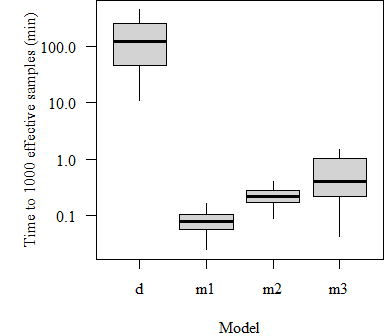

***Figure S2****. Boxplot of the time required to reach 1000 effective samples of the least converged parameter for discrete (d) and marginalized (m1, m2, and m3) multistate Jolly Seber models applied to 100 complete simulated data sets.*

R and JAGS code for writing model files, simulating data, running models, and extracting results, are as follows:

```
#----------------------------------------------------------------------------# Write models to files
# 1. Discrete latent state model (d)
{
sink("js-ms-d.txt")
cat("
model {

#--------------------------------------
# phi: survival probability
# gamma: removal entry probability
# p: capture probability
#--------------------------------------

# Priors and constraints
for (t in 1:(n.occasions-1)){
   phi[t] <- mean.phi     # Constant survival
   gamma[t] ~ dunif(0,1)  # Time-dependent entry
   p[t] <- mean.p         # Constant capture probability
   }

mean.phi ~ dunif(0,1)    # Prior for mean survival
mean.p ~ dunif(0,1)      # Prior for mean capture

#--------------------------------------
# States (S):
# 1 not yet entered
# 2 alive
# 3 dead
#--------------------------------------

# Probabilities of state S(t+1) given S(t)
   for (t in 1:(n.occasions-1)){
      omega[1,1,t] <- 1-gamma[t]
      omega[1,2,t] <- gamma[t]
      omega[1,3,t] <- 0
      omega[2,1,t] <- 0
      omega[2,2,t] <- phi[t]
      omega[2,3,t] <- 1-phi[t]
      omega[3,1,t] <- 0
      omega[3,2,t] <- 0
      omega[3,3,t] <- 1

#--------------------------------------
# Observations (O):
# 1 captured
# 2 not captured
#--------------------------------------

# Probabilities of O(t) given S(t)
      rho[1,1,t] <- 0
      rho[1,2,t] <- 1
      rho[2,1,t] <- p[t]
      rho[2,2,t] <- 1-p[t]
      rho[3,1,t] <- 0
      rho[3,2,t] <- 1
      } #t

# Likelihood
for (i in 1:M){
   # Define latent state at first occasion
   s[i,1] ~ dcat(1)   # Make sure that all M individuals are in state 1 at t=1
   for (t in 2:n.occasions){
      # State process: draw S(t) given S(t-1)
      s[i,t] ~ dcat(omega[s[i,t-1],,t-1])
      # Observation process: draw O(t) given S(t)
      y[i,t] ~ dcat(rho[s[i,t],,t-1])
      } #t
   } #i

# Derived population parameters
for (t in 1:(n.occasions-1)){
   qgamma[t] <- 1-gamma[t]
   }
c[1] <- gamma[1]
for (t in 2:(n.occasions-1)){
   c[t] <- gamma[t] * prod(qgamma[1:(t-1)])
   } # t
psi <- sum(c[1:(n.occasions-1)])            # inclusion probability
for (t in 1:(n.occasions-1)){
   b[t] <- c[t] / psi      # entry probability
   } # t

for (i in 1:M){
   for (t in 2:n.occasions){
      al[i,t-1] <- equals(s[i,t],2)
      } # t
   for (t in 1:(n.occasions-1)){
      d[i,t] <- equals(s[i,t]-al[i,t],0)
      } #t
   alive[i] <- sum(al[i,])
   } # i

for (t in 1:(n.occasions-1)){
   N[t] <- sum(al[,t])        # population size
   B[t] <- sum(d[,t])         # number of entries
   } # t
for (i in 1:M){
   w[i] <- 1-equals(alive[i],0)
   } # i
for(t in 1:(n.occasions-2)){
  lambda[t] <- N[t+1]/N[t] # population growth rate
  f1[t] <- B[t+1]/N[t] # recruitment rate
  f2[t] <- lambda[t] - phi[t]
} # t
Nsuper <- sum(w[])            # superpopulation size
}
",fill = TRUE)
sink()


# 2. Marginalized multistate Jolly Seber model - derived parameters (m1)

sink("js-ms-m1.txt")
cat("

model{

  # Priors and constraints
  for (t in 1:(n.occasions-1)){
    phi[t] <- mean.phi     # Constant survival
    gamma[t] ~ dunif(0,1)  # Time-dependent entry probabilities
    p[t] <- mean.p         # Constant capture probability
  }

  mean.phi ~ dunif(0,1)    # Prior for mean survival
  mean.p ~ dunif(0,1)      # Prior for mean capture

  for (t in 1:(n.occasions-1)){

  #--------------------------------------------------------
  # states (S)
  # 1 = not yet entered
  # 2 = alive
  # 3 = dead
  #--------------------------------------------------------

  # Probabilities of state S(t+1) given S(t)
    omega[1,1,t] <- 1-gamma[t]
    omega[1,2,t] <- gamma[t]
    omega[1,3,t] <- 0
    omega[2,1,t] <- 0
    omega[2,2,t] <- phi[t]
    omega[2,3,t] <- 1-phi[t]
    omega[3,1,t] <- 0
    omega[3,2,t] <- 0
    omega[3,3,t] <- 1

  #--------------------------------------------------------
  # observations (O)
  # 1 = captured
  # 2 = not captured
  #--------------------------------------------------------

  # Probabilities of O(t) given S(t)
    rho[1,1,t] <- 0
    rho[1,2,t] <- 1
    rho[2,1,t] <- p[t]
    rho[2,2,t] <- 1 - p[t]
    rho[3,1,t] <- 0
    rho[3,2,t] <- 1

  } # t

  # Likelihood
  for (i in 1:MS){
    # Define latent state at first occasion
    zeta[i,1,1] <- 1   # Set all individuals to state 1 (not yet entered) at t=1
    zeta[i,1,2] <- 0
    zeta[i,1,3] <- 0

    for (t in 2:(n.occasions)){
      # State process: draw S(t+1) given S(t)
      zeta[i,t,1] <- inprod(zeta[i,(t-1),1:3], omega[1:3,1,(t-1)] *
      rho[1,ys[i,t],(t-1)])
      zeta[i,t,2] <- inprod(zeta[i,(t-1),1:3], omega[1:3,2,(t-1)] *
      rho[2,ys[i,t],(t-1)])
      zeta[i,t,3] <- inprod(zeta[i,(t-1),1:3], omega[1:3,3,(t-1)] *
      rho[3,ys[i,t],(t-1)])
    } # t
    lik[i] <- sum(zeta[i,(n.occasions),1:3])
    fr[i] ~ dbin(lik[i], FR[i])
  } # i

  # Calculate derived population parameters
  for (t in 1:(n.occasions-1)){
    qgamma[t] <- 1-gamma[t]
  } # t
  c[1] <- gamma[1]
  N[1] <- B[1]
  for (t in 2:(n.occasions-1)){
    c[t] <- gamma[t] * prod(qgamma[1:(t-1)])
    N[t] <- B[t] + phi[t] * N[t-1]
  } # t
  psi <- sum(c[1:(n.occasions-1)])    # inclusion probability
  for (t in 1:(n.occasions-1)){
    b[t] <- c[t] / psi      # entry probability
    B[t] <- c[t]*M       # number of entries
  } # t

  for(t in 1:(n.occasions-2)){
  lambda[t] <- N[t+1]/N[t] # population growth rate
  f1[t] <- B[t+1]/N[t] # recruitment rate
  f2[t] <- lambda[t] - phi[t]
  } # t

  Nsuper <- sum(B[1:(n.occasions-1)]) # superpopulation size
}

",fill=TRUE)
sink()

# 3. Marginalized multistate Jolly Seber model - simulated N and B (m2)

sink("js-ms-m2.txt")
cat("
model{

  # Priors and constraints
  for (t in 1:(n.occasions-1)){
    phi[t] <- mean.phi     # Constant survival
    gamma[t] ~ dunif(0,1)  # Time-dependent entry probabilities
    p[t] <- mean.p         # Constant capture probability
  }

  mean.phi ~ dunif(0,1)    # Prior for mean survival
  mean.p ~ dunif(0,1)      # Prior for mean capture

  for (t in 1:(n.occasions-1)){

  #--------------------------------------------------------
  # states (S)
  # 1 = not yet entered
  # 2 = alive
  # 3 = dead
  #--------------------------------------------------------

  # Probabilities of state S(t+1) given S(t)
    omega[1,1,t] <- 1-gamma[t]
    omega[1,2,t] <- gamma[t]
    omega[1,3,t] <- 0
    omega[2,1,t] <- 0
    omega[2,2,t] <- phi[t]
    omega[2,3,t] <- 1-phi[t]
    omega[3,1,t] <- 0
    omega[3,2,t] <- 0
    omega[3,3,t] <- 1

  #--------------------------------------------------------
  # observations (O)
  # 1 = captured
  # 2 = not captured
  #--------------------------------------------------------

  # Probabilities of O(t) given S(t)
    rho[1,1,t] <- 0
    rho[1,2,t] <- 1
    rho[2,1,t] <- p[t]
    rho[2,2,t] <- 1 - p[t]
    rho[3,1,t] <- 0
    rho[3,2,t] <- 1

  } # t

  # Likelihood
  for (i in 1:MS){
    # Define latent state at first occasion
    zeta[i,1,1] <- 1   # Set all individuals to state 1 (not yet entered) at t=1
    zeta[i,1,2] <- 0
    zeta[i,1,3] <- 0

    for (t in 2:(n.occasions)){
      # State process: draw S(t+1) given S(t)
      zeta[i,t,1] <- inprod(zeta[i,(t-1),1:3], omega[1:3,1,(t-1)] *
      rho[1,ys[i,t],(t-1)])
      zeta[i,t,2] <- inprod(zeta[i,(t-1),1:3], omega[1:3,2,(t-1)] *
      rho[2,ys[i,t],(t-1)])
      zeta[i,t,3] <- inprod(zeta[i,(t-1),1:3], omega[1:3,3,(t-1)] *
      rho[3,ys[i,t],(t-1)])
    } # t
    lik[i] <- sum(zeta[i,(n.occasions),1:3])
    fr[i] ~ dbin(lik[i], FR[i])
  } # i

  # Calculate derived population parameters
  for (t in 1:(n.occasions-1)){
    qgamma[t] <- 1-gamma[t]
  } # t
  c[1] <- gamma[1]
  for (t in 2:(n.occasions-1)){
    c[t] <- gamma[t] * prod(qgamma[1:(t-1)])
  } # t
  psi <- sum(c[1:(n.occasions-1)])    # inclusion probability
  for (t in 1:(n.occasions-1)){
    b[t] <- c[t] / psi                # entry probability
  } # t

  p0_inc<-sum(zeta[MS,n.occasions,2:3])/sum(zeta[MS,n.occasions,1:3])
  U ~ dbin(p0_inc,FR[MS]) #assumes empty CH is always the last one
  Nsuper <- sum(FR[1:(MS-1)])+U # superpopulation size

# This loop calculates the likelihood that an individual recruits during time
# step poss[i,1], dies during time step poss[i,2] and is never seen while alive

for (i in 1:nposs){
    lik0[i]<-omega[1,2,poss[i,1]]*((1-equals(poss[i,2],(n.occasions-1)))*
    (prod(rho[2,2,poss[i,1]:poss[i,2]])*prod(omega[2,2,poss[i,1]:(n.occasions-1)])*
    (omega[2,3,poss[i,2]])/prod(omega[2,2,(poss[i,2]):(n.occasions-1)]))+
        (equals(poss[i,2],(n.occasions-1)))*(prod(rho[2,2,poss[i,1]:poss[i,2]])
        *prod(omega[2,2,poss[i,1]:(n.occasions-1)])/prod(omega[2,2,poss[i,2]:(n.occasions-1)])))
}

# The next set of loops determines for each individual (linked to summarized
# capture history by lookup) determine if an underlying set of underlying dynamics
# (i.e., recruit and death period) is possible given their capture history

for (i in 1:Nobs_ind){
    for (j in 1:(n.occasions-1)){
        for (k in 1:nposs){
            inc_a[i,j,k]<-equals(poss[k,1],j)*equals(poss_obs[k,lookup[i]],1)
            inc_l[i,j,k]<-equals(poss[k,2],j)*equals(poss_obs[k,lookup[i]],1)
            }
        p_a[i,j]<-inprod(inc_a[i,j,1:nposs],lik0[1:nposs])
        p_l[i,j]<-inprod(inc_l[i,j,1:nposs],lik0[1:nposs])
        }
    ar[i]~dcat(p_a[i,])
    li[i]~dcat(p_l[i,])
    A[i]<-ar[i]
    L[i]<-li[i]
    for (j in 1:(n.occasions-1)){
        S[i,j]<-step(j-A[i])*step(L[i]-j)
        }
}

# The next set of loops determines the likelihood of an underlying set of
# underlying dynamics (i.e., recruit and death period) given an individual
# was never seen.

for (j in 1:(n.occasions-1)){
    for (k in 1:nposs){
        inc_a_no[j,k]<-equals(poss[k,1],j)
        inc_l_no[j,k]<-equals(poss[k,2],j)
        }
    p_a_no[j]<-inprod(inc_a_no[j,1:nposs],lik0[1:nposs])
    p_l_no[j]<-inprod(inc_l_no[j,1:nposs],lik0[1:nposs])
    }

for (i in (Nobs_ind+1):(Npos_ind)){
    ar[i]~dcat(p_a_no)
    li[i]~dcat(p_l_no)
    A[i]<-ar[i]*step(Nobs_ind+U-i)
    L[i]<-li[i]*step(Nobs_ind+U-i)
    for (j in 1:(n.occasions-1)){
        S[i,j]<-step(j-A[i])*step(L[i]-j)
        }
    }

for (t in 1:(n.occasions-1)){
  B[t] <- sum(equals(A[1:(Npos_ind)],t)) # number of entries
  N[t] <- sum(S[1:(Npos_ind),t])
  } # t
  for(t in 1:(n.occasions-2)){
  lambda[t] <- N[t+1]/N[t] # population growth rate
  f1[t] <- B[t+1]/N[t]# recruitment rate
  f2[t] <- lambda[t] - phi[t]
  } # t
 }
    ", fill =TRUE)
sink()

# 3. Marginalized multistate Jolly Seber model - simulated N, not B (m3)

sink("js-ms-m3.txt")
cat("
model{

  # Priors and constraints
  for (t in 1:(n.occasions-1)){
    phi[t] <- mean.phi     # Constant survival
    gamma[t] ~ dbeta(1/(n.occasions-1), 2-t/(n.occasions-1)) # Time-dependent entry probabilities
    p[t] <- mean.p         # Constant capture probability
  }

  mean.phi ~ dunif(0,1)    # Prior for mean survival
  mean.p ~ dunif(0,1)      # Prior for mean capture

  for (t in 1:(n.occasions-1)){

  #--------------------------------------------------------
  # states (S)
  # 1 = not yet entered
  # 2 = alive
  # 3 = dead
  #--------------------------------------------------------

  # Probabilities of state S(t+1) given S(t)
    omega[1,1,t] <- 1-gamma[t]
    omega[1,2,t] <- gamma[t]
    omega[1,3,t] <- 0
    omega[2,1,t] <- 0
    omega[2,2,t] <- phi[t]
    omega[2,3,t] <- 1-phi[t]
    omega[3,1,t] <- 0
    omega[3,2,t] <- 0
    omega[3,3,t] <- 1

  #--------------------------------------------------------
  # observations (O)
  # 1 = captured
  # 2 = not captured
  #--------------------------------------------------------

  # Probabilities of O(t) given S(t)
    rho[1,1,t] <- 0
    rho[1,2,t] <- 1
    rho[2,1,t] <- p[t]
    rho[2,2,t] <- 1 - p[t]
    rho[3,1,t] <- 0
    rho[3,2,t] <- 1

  } # t

  # Likelihood
  for (i in 1:MS){
    # Define latent state at first occasion
    zeta[i,1,1] <- 1   # Set all individuals to state 1 (not yet entered) at t=1
    zeta[i,1,2] <- 0
    zeta[i,1,3] <- 0

    for (t in 2:(n.occasions)){
      zeta[i,t,1] <- inprod(zeta[i,(t-1),1:3], omega[1:3,1,(t-1)] *
      rho[1,ys[i,t],(t-1)])
      zeta[i,t,2] <- inprod(zeta[i,(t-1),1:3], omega[1:3,2,(t-1)] *
      rho[2,ys[i,t],(t-1)])
      zeta[i,t,3] <- inprod(zeta[i,(t-1),1:3], omega[1:3,3,(t-1)] *
      rho[3,ys[i,t],(t-1)])
    } # t
    lik[i] <- sum(zeta[i,(n.occasions),1:3])
    fr[i] ~ dbin(lik[i], FR[i])
  } # i

  # Calculate derived population parameters
  for (t in 1:(n.occasions-1)){
    qgamma[t] <- 1-gamma[t]
  } # t
  c[1] <- gamma[1]
  for (t in 2:(n.occasions-1)){
    c[t] <- gamma[t] * prod(qgamma[1:(t-1)])
  } # t
  psi <- sum(c[1:(n.occasions-1)])    # inclusion probability
  for (t in 1:(n.occasions-1)){
    b[t] <- c[t] / psi                # entry probability
    B[t] <- c[t]*M
  } # t

  p0_inc<-sum(zeta[MS,n.occasions,2:3])/sum(zeta[MS,n.occasions,1:3])
  U ~ dbin(p0_inc,FR[MS]) #assumes empty CH is always the last one
  Nsuper <- sum(FR[1:(MS-1)])+U # superpopulation size

  for (i in 1:M){
  # Simulations need to be done for individuals even though zeta can be calculated
  # for summarized capture histories. lookup refers each individual in M, to the
  # appropriate summarized capture history.
  for (t in 2:(n.occasions)){
  al[i,t-1] ~ dbern(zeta[lookup[i],t,2]/sum(zeta[lookup[i],t,1:3]))
  } # t
  } # i

  for (t in 1:(n.occasions-1)){
  N[t] <- sum(al[1:M,t]) # population size
  } # t

  for(t in 1:(n.occasions-2)){
  lambda[t] <- N[t+1]/N[t] # population growth rate

  f1[t] <- B[t+1]/N[t]# recruitment rate
  f2[t] <- lambda[t] - phi[t] # recruitment rate
  } # t
 }
    ", fill =TRUE)
sink()
}

# some functions ------------------------------------------------------------

# Simulate capture histories (from Kery and Schaub 2012)

simul.js <- function(PHI, P, b, Nsuper){
  B <- rmultinom(1, Nsuper, b) # Generate no. of entering ind. per occasion
  n.occasions <- dim(PHI)[2] + 1
  CH.sur <- CH.p <- matrix(0, ncol = n.occasions, nrow = Nsuper)
  # Define a vector with the occasion of entering the population
  ent.occ <- numeric()
  for (t in 1:n.occasions){
    ent.occ <- c(ent.occ, rep(t, B[t]))
  }
  # Simulating survival
  for (i in 1:Nsuper){
    CH.sur[i, ent.occ[i]] <- 1   # Write 1 when ind. enters the pop.
    if (ent.occ[i] == n.occasions) next
    for (t in (ent.occ[i]+1):n.occasions){
      # Bernoulli trial: has individual survived occasion?
      sur <- rbinom(1, 1, PHI[i,t-1])
      ifelse (sur==1, CH.sur[i,t] <- 1, break)
    } #t
  } #i
  # Simulating capture
  for (i in 1:Nsuper){
    CH.p[i,] <- rbinom(n.occasions, 1, P[i,])
  } #i
  # Full capture-recapture matrix
  CH <- CH.sur * CH.p

  # Remove individuals never captured
  cap.sum <- rowSums(CH)
  never <- which(cap.sum == 0)
  CH <- CH[-never,]
  Nt <- colSums(CH.sur)    # Actual population size
  return(list(CH=CH, B=B, Nsuper=Nt))
}


# Collapse capture histories (from Yackulic et al. 2020)
collapse.ch <- function(ch){
  ch.char = apply(ch, 1, function(x) paste(x, collapse = ","))

  ch.sum.out = t(sapply(strsplit(names(table(ch.char)), split = ","),
                        as.numeric))
  fr.out = as.numeric(as.vector(table(ch.char)))

  return(list(ch.sum.out, fr.out))
}

#----------------------------------------------------------------------------

library(jagsUI)
library(dplyr)
##
## Attaching package: 'dplyr'
## The following objects are masked from 'package:stats':
##
##     filter, lag
## The following objects are masked from 'package:base':
##
##     intersect, setdiff, setequal, union
library(data.table)
##
## Attaching package: 'data.table'
## The following objects are masked from 'package:dplyr':
##
##     between, first, last
# some functions ---------------------------------------------------------------

# Simulate capture histories (from Kery and Schaub 2012)

simul.js <- function(PHI, P, b, Nsuper){
  B <- rmultinom(1, Nsuper, b) # Generate no. of entering ind. per occasion
  n.occasions <- dim(PHI)[2] + 1
  CH.sur <- CH.p <- matrix(0, ncol = n.occasions, nrow = Nsuper)
  # Define a vector with the occasion of entering the population
  ent.occ <- numeric()
  for (t in 1:n.occasions){
    ent.occ <- c(ent.occ, rep(t, B[t]))
  }
  # Simulating survival
  for (i in 1:Nsuper){
    CH.sur[i, ent.occ[i]] <- 1   # Write 1 when ind. enters the pop.
    if (ent.occ[i] == n.occasions) next
    for (t in (ent.occ[i]+1):n.occasions){
      # Bernoulli trial: has individual survived occasion?
      sur <- rbinom(1, 1, PHI[i,t-1])
      ifelse (sur==1, CH.sur[i,t] <- 1, break)
    } #t
  } #i
  # Simulating capture
  for (i in 1:Nsuper){
    CH.p[i,] <- rbinom(n.occasions, 1, P[i,])
  } #i
  # Full capture-recapture matrix
  CH <- CH.sur * CH.p

  # Remove individuals never captured
  cap.sum <- rowSums(CH)
  never <- which(cap.sum == 0)
  CH <- CH[-never,]
  Nt <- colSums(CH.sur)    # Actual population size
  return(list(CH=CH, B=B, Nsuper=Nt))
}


# Fill in initial values for s. Set to 1 (not entered) for all occasions prior to
# first capture, 2 for occasions from first to last capture, and 3 for occasions
# after last capture (from Kery and Schaub 2012).

js.multistate.init <- function(ch, ns){
  ch[ch==2] <- NA
  state <- ch
  for (i in 1:nrow(ch)){
    n1 <- min(which(ch[i,]==1))
    n2 <- max(which(ch[i,]==1))
    state[i,n1:n2] <- 2
  }
  state[state==0] <- NA
  get.first <- function(x) min(which(!is.na(x)))
  get.last <- function(x) max(which(!is.na(x)))
  f <- apply(state, 1, get.first)
  l <- apply(state, 1, get.last)
  for (i in 1:nrow(ch)){
    state[i,1:f[i]] <- 1
    if(l[i]!=ncol(ch)) state[i, (l[i]+1):ncol(ch)] <- 3
    state[i, f[i]] <- 2
  }
  state <- rbind(state, matrix(1, ncol = ncol(ch), nrow = ns))
  return(state)
}

# Collapse capture histories (from Yackulic et al. 2020)

collapse.ch <- function(ch){
  options(warn=-1)
  ch.char = apply(ch, 1, function(x) paste(x, collapse = ","))
  ch.sum.out = t(sapply(strsplit(names(table(ch.char)), split = ","), as.numeric))
  fr.out = as.numeric(as.vector(table(ch.char)))
  return(list(ch.sum.out, fr.out))
}

nrep <- 100
# Define parameter values
n.occasions <- 7                         # Number of capture occasions
Nsuper <- 400                            # Superpopulation size
phi <- rep(0.7, n.occasions-1)           # Survival probabilities
b <- c(0.40, rep(0.11, n.occasions-1))   # Entry probabilities
p <- rep(0.5, n.occasions)               # Capture probabilities

PHI <- matrix(rep(phi, (n.occasions-1)*Nsuper), ncol = n.occasions-1, nrow =
                Nsuper, byrow = TRUE)
P <- matrix(rep(p, n.occasions*Nsuper), ncol = n.occasions, nrow = Nsuper,
            byrow = TRUE)

# Create data frames to store results
js.med.df <- data.frame(model=character(), Nsuper=numeric(), phi=numeric(),
                       b1=numeric(), b2=numeric(), b3=numeric(), b4=numeric(),
                        b5=numeric(), b6=numeric(), b7=numeric(), p=numeric(),
                        f1=numeric(), f2=numeric(), f3=numeric(), f4=numeric(),
                        f5=numeric(), f6=numeric(), N1=numeric(), N2=numeric(),
                        N3=numeric(), N4=numeric(), N5=numeric(), N6=numeric(),
                        N7=numeric(), lambda1=numeric(), lambda2=numeric(),
                        lambda3=numeric(),lambda4=numeric(),lambda5=numeric(),
                        lambda6=numeric())

js.sd.df <- data.frame(model=character(), Nsuper=numeric(), phi=numeric(),
                       b1=numeric(), b2=numeric(), b3=numeric(), b4=numeric(),
                       b5=numeric(), b6=numeric(), b7=numeric(), p=numeric(),
                       f1=numeric(), f2=numeric(), f3=numeric(), f4=numeric(),
                       f5=numeric(), f6=numeric(), N1=numeric(), N2=numeric(),
                       N3=numeric(), N4=numeric(), N5=numeric(), N6=numeric(),
                       N7=numeric(), lambda1=numeric(), lambda2=numeric(),
                       lambda3=numeric(),lambda4=numeric(),lambda5=numeric(),
                       lambda6=numeric())

js.025.df <- data.frame(model=character(), Nsuper=numeric(), phi=numeric(),
                        b1=numeric(), b2=numeric(), b3=numeric(), b4=numeric(),
                        b5=numeric(), b6=numeric(), b7=numeric(), p=numeric(),
                        f1=numeric(), f2=numeric(), f3=numeric(), f4=numeric(),
                        f5=numeric(), f6=numeric(), N1=numeric(), N2=numeric(),
                        N3=numeric(), N4=numeric(), N5=numeric(), N6=numeric(),
                        N7=numeric(), lambda1=numeric(), lambda2=numeric(),
                        lambda3=numeric(),lambda4=numeric(),lambda5=numeric(),
                        lambda6=numeric())

js.975.df <- data.frame(model=character(), Nsuper=numeric(), phi=numeric(),
                        b1=numeric(), b2=numeric(), b3=numeric(), b4=numeric(),
                        b5=numeric(), b6=numeric(), b7=numeric(), p=numeric(),
                        f1=numeric(), f2=numeric(), f3=numeric(), f4=numeric(),
                        f5=numeric(), f6=numeric(), N1=numeric(), N2=numeric(),
                        N3=numeric(), N4=numeric(), N5=numeric(), N6=numeric(),
                        N7=numeric(), lambda1=numeric(), lambda2=numeric(),
                        lambda3=numeric(),lambda4=numeric(),lambda5=numeric(),
                        lambda6=numeric())

js.mcmc.df <- data.frame(model=character(), rt=numeric(), min.ess=numeric(),
                         max.ess=numeric(), med.ess=numeric(), max.Rhat=numeric())

# Simulate data and run models

# define number of iterations
nrep <- 100

js <- function(nrep){

  for (i in 1:nrep){

    # Simulate capture histories
    sim <- simul.js(PHI, P, b, Nsuper)
    CH <- sim$CH
    # Add dummy occasion
    CH.du <- cbind(rep(0, dim(CH)[1]), CH)
    # Augment data
    ns <- 500
    CH.ms <- rbind(CH.du, matrix(0, ncol = dim(CH.du)[2], nrow = ns))
    # Recode CH matrix: 2 = Not captured , 1 = captured
    CH.ms[CH.ms==0] <- 2
    # collapse to unique capture histories and frequencies
    chsum <- collapse.ch(CH.ms)
    ys <- chsum[[1]]
    FR <- chsum[[2]]
    M <- sum(FR)

    # MCMC settings
    ni <- 20000
    nt <- 3
    nb <- 5000
    nc <- 3

    # Parameters to monitor
    parameters <- c("Nsuper", "mean.phi", "b", "mean.p", "N", "lambda",
                    "f1", "f2")

    # model d -----------------------------------------------------------

    # Data for discrete multistate JS model
    jags.data <- list(y = CH.ms, n.occasions = dim(CH.ms)[2], M = dim(CH.ms)[1])
    # initial values for s
    sst <- js.multistate.init(CH.du, ns)
    # set initial values
    inits <- function(){list(mean.phi = runif(1, .6, .8),
                             mean.p = runif(1, .4, .6), s = sst)}

    # run model
    js.ms.d <- jags(jags.data, inits, parameters, "js-ms-d.txt", n.chains = nc,
                    n.thin = nt, n.iter = ni, n.burnin = nb, parallel=TRUE)

    # extract results

    js.med.df <- rbind(js.med.df, data.frame(model = "d",
                Nsuper = median(js.ms.d$sims.list$Nsuper),
                phi = median(js.ms.d$sims.list$mean.phi),
                b1 = median(js.ms.d$sims.list$b[,1]),
                b2 = median(js.ms.d$sims.list$b[,2]),
                b3 = median(js.ms.d$sims.list$b[,3]),
                b4 = median(js.ms.d$sims.list$b[,4]),
                b5 = median(js.ms.d$sims.list$b[,5]),
                b6 = median(js.ms.d$sims.list$b[,6]),
                b7 = median(js.ms.d$sims.list$b[,7]),
                p = median(js.ms.d$sims.list$mean.p),
                f1 = median(js.ms.d$sims.list$f1[,1]),
                f2 = median(js.ms.d$sims.list$f1[,2]),
                f3 = median(js.ms.d$sims.list$f1[,3]),
                f4 = median(js.ms.d$sims.list$f1[,4]),
                f5 = median(js.ms.d$sims.list$f1[,5]),
                f6 = median(js.ms.d$sims.list$f1[,6]),
                N1 = median(js.ms.d$sims.list$N[,1]),
                N2 = median(js.ms.d$sims.list$N[,2]),
                N3 = median(js.ms.d$sims.list$N[,3]),
                N4 = median(js.ms.d$sims.list$N[,4]),
                N5 = median(js.ms.d$sims.list$N[,5]),
                N6 = median(js.ms.d$sims.list$N[,6]),
                N7 = median(js.ms.d$sims.list$N[,7]),
                lambda1 = median(js.ms.d$sims.list$lambda[,1]),
                lambda2 = median(js.ms.d$sims.list$lambda[,2]),
                lambda3 = median(js.ms.d$sims.list$lambda[,3]),
                lambda4 = median(js.ms.d$sims.list$lambda[,4]),
                lambda5 = median(js.ms.d$sims.list$lambda[,5]),
                lambda6 = median(js.ms.d$sims.list$lambda[,6])))

    js.sd.df <- rbind(js.sd.df, data.frame(model = "d",
                Nsuper = sd(js.ms.d$sims.list$Nsuper),
                phi = sd(js.ms.d$sims.list$mean.phi),
                b1 = sd(js.ms.d$sims.list$b[,1]),
                b2 = sd(js.ms.d$sims.list$b[,2]),
                b3 = sd(js.ms.d$sims.list$b[,3]),
                b4 = sd(js.ms.d$sims.list$b[,4]),
                b5 = sd(js.ms.d$sims.list$b[,5]),
                b6 = sd(js.ms.d$sims.list$b[,6]),
                b7 = sd(js.ms.d$sims.list$b[,7]),
                p = sd(js.ms.d$sims.list$mean.p),
                f1 = sd(js.ms.d$sims.list$f1[,1]),
                f2 = sd(js.ms.d$sims.list$f1[,2]),
                f3 = sd(js.ms.d$sims.list$f1[,3]),
                f4 = sd(js.ms.d$sims.list$f1[,4]),
                f5 = sd(js.ms.d$sims.list$f1[,5]),
                f6 = sd(js.ms.d$sims.list$f1[,6]),
                N1 = sd(js.ms.d$sims.list$N[,1]),
                N2 = sd(js.ms.d$sims.list$N[,2]),
                N3 = sd(js.ms.d$sims.list$N[,3]),
                N4 = sd(js.ms.d$sims.list$N[,4]),
                N5 = sd(js.ms.d$sims.list$N[,5]),
                N6 = sd(js.ms.d$sims.list$N[,6]),
                N7 = sd(js.ms.d$sims.list$N[,7]),
                lambda1 = sd(js.ms.d$sims.list$lambda[,1]),
                lambda2 = sd(js.ms.d$sims.list$lambda[,2]),
                lambda3 = sd(js.ms.d$sims.list$lambda[,3]),
                lambda4 = sd(js.ms.d$sims.list$lambda[,4]),
                lambda5 = sd(js.ms.d$sims.list$lambda[,5]),
                lambda6 = sd(js.ms.d$sims.list$lambda[,6])))

    js.025.df <- rbind(js.025.df, data.frame(model = "d",
                Nsuper = quantile(js.ms.d$sims.list$Nsuper, probs = 0.025),
                phi = quantile(js.ms.d$sims.list$mean.phi, probs = 0.025),
                b1 = quantile(js.ms.d$sims.list$b[,1], probs = 0.025),
                b2 = quantile(js.ms.d$sims.list$b[,2], probs = 0.025),
                b3 = quantile(js.ms.d$sims.list$b[,3], probs = 0.025),
                b4 = quantile(js.ms.d$sims.list$b[,4], probs = 0.025),
                b5 = quantile(js.ms.d$sims.list$b[,5], probs = 0.025),
                b6 = quantile(js.ms.d$sims.list$b[,6], probs = 0.025),
                b7 = quantile(js.ms.d$sims.list$b[,7], probs = 0.025),
                p = quantile(js.ms.d$sims.list$mean.p, probs = 0.025),
                f1 = quantile(js.ms.d$sims.list$f1[,1], probs = 0.025),
                f2 = quantile(js.ms.d$sims.list$f1[,2], probs = 0.025),
                f3 = quantile(js.ms.d$sims.list$f1[,3], probs = 0.025),
                f4 = quantile(js.ms.d$sims.list$f1[,4], probs = 0.025),
                f5 = quantile(js.ms.d$sims.list$f1[,5], probs = 0.025),
                f6 = quantile(js.ms.d$sims.list$f1[,6], probs = 0.025),
                N1 = quantile(js.ms.d$sims.list$N[,1], probs = 0.025),
                N2 = quantile(js.ms.d$sims.list$N[,2], probs = 0.025),
                N3 = quantile(js.ms.d$sims.list$N[,3], probs = 0.025),
                N4 = quantile(js.ms.d$sims.list$N[,4], probs = 0.025),
                N5 = quantile(js.ms.d$sims.list$N[,5], probs = 0.025),
                N6 = quantile(js.ms.d$sims.list$N[,6], probs = 0.025),
                N7 = quantile(js.ms.d$sims.list$N[,7], probs = 0.025),
                lambda1 = quantile(js.ms.d$sims.list$lambda[,1], probs = 0.025),
                lambda2 = quantile(js.ms.d$sims.list$lambda[,2], probs = 0.025),
                lambda3 = quantile(js.ms.d$sims.list$lambda[,3], probs = 0.025),
                lambda4 = quantile(js.ms.d$sims.list$lambda[,4], probs = 0.025),
                lambda5 = quantile(js.ms.d$sims.list$lambda[,5], probs = 0.025),
                lambda6 = quantile(js.ms.d$sims.list$lambda[,6], probs = 0.025)))

    js.975.df <- rbind(js.975.df, data.frame(model = "d",
                Nsuper = quantile(js.ms.d$sims.list$Nsuper, probs = 0.975),
                phi = quantile(js.ms.d$sims.list$mean.phi, probs = 0.975),
                b1 = quantile(js.ms.d$sims.list$b[,1], probs = 0.975),
                b2 = quantile(js.ms.d$sims.list$b[,2], probs = 0.975),
                b3 = quantile(js.ms.d$sims.list$b[,3], probs = 0.975),
                b4 = quantile(js.ms.d$sims.list$b[,4], probs = 0.975),
                b5 = quantile(js.ms.d$sims.list$b[,5], probs = 0.975),
                b6 = quantile(js.ms.d$sims.list$b[,6], probs = 0.975),
                b7 = quantile(js.ms.d$sims.list$b[,7], probs = 0.975),
                p = quantile(js.ms.d$sims.list$mean.p, probs = 0.975),
                f1 = quantile(js.ms.d$sims.list$f1[,1], probs = 0.975),
                f2 = quantile(js.ms.d$sims.list$f1[,2], probs = 0.975),
                f3 = quantile(js.ms.d$sims.list$f1[,3], probs = 0.975),
                f4 = quantile(js.ms.d$sims.list$f1[,4], probs = 0.975),
                f5 = quantile(js.ms.d$sims.list$f1[,5], probs = 0.975),
                f6 = quantile(js.ms.d$sims.list$f1[,6], probs = 0.975),
                N1 = quantile(js.ms.d$sims.list$N[,1], probs = 0.975),
                N2 = quantile(js.ms.d$sims.list$N[,2], probs = 0.975),
                N3 = quantile(js.ms.d$sims.list$N[,3], probs = 0.975),
                N4 = quantile(js.ms.d$sims.list$N[,4], probs = 0.975),
                N5 = quantile(js.ms.d$sims.list$N[,5], probs = 0.975),
                N6 = quantile(js.ms.d$sims.list$N[,6], probs = 0.975),
                N7 = quantile(js.ms.d$sims.list$N[,7], probs = 0.975),
                lambda1 = quantile(js.ms.d$sims.list$lambda[,1], probs = 0.975),
                lambda2 = quantile(js.ms.d$sims.list$lambda[,2], probs = 0.975),
                lambda3 = quantile(js.ms.d$sims.list$lambda[,3], probs = 0.975),
                lambda4 = quantile(js.ms.d$sims.list$lambda[,4], probs = 0.975),
                lambda5 = quantile(js.ms.d$sims.list$lambda[,5], probs = 0.975),
                lambda6 = quantile(js.ms.d$sims.list$lambda[,6], probs = 0.975)))

    js.mcmc.df <- rbind(js.mcmc.df, data.frame(model="d",
                  rt=js.ms.d$mcmc.info$elapsed.mins,
                  min.ess=min(unlist(js.ms.d$n.eff)),
                  max.ess=max(unlist(js.ms.d$n.eff)),
                  med.ess=median(unlist(js.ms.d$n.eff)),
                  max.Rhat=max(unlist(js.ms.d$Rhat))))

    # model m1 -------------------------------------------------------

    # data for marginalized JS model
    jags.data<- list(FR=FR, fr=FR, ys = ys, n.occasions=dim(ys)[2], MS=dim(ys)[1],
                     M = M)
    # set initial values
    inits <- function(){ list(mean.phi = .45, mean.p = runif(1, .4, .6))}
    # run model
    js.ms.m1 <- jags(jags.data, inits, parameters, "js-ms-m1.txt", n.chains = nc,
                    n.thin =nt, n.iter = ni, n.burnin = nb, parallel=TRUE)

    # extract results
    js.med.df <- rbind(js.med.df, data.frame(model = "m1",
                 Nsuper = median(js.ms.m1$sims.list$Nsuper),
                 phi = median(js.ms.m1$sims.list$mean.phi),
                 b1 = median(js.ms.m1$sims.list$b[,1]),
                 b2 = median(js.ms.m1$sims.list$b[,2]),
                 b3 = median(js.ms.m1$sims.list$b[,3]),
                 b4 = median(js.ms.m1$sims.list$b[,4]),
                 b5 = median(js.ms.m1$sims.list$b[,5]),
                 b6 = median(js.ms.m1$sims.list$b[,6]),
                 b7 = median(js.ms.m1$sims.list$b[,7]),
                 p = median(js.ms.m1$sims.list$mean.p),
                 f1 = median(js.ms.m1$sims.list$f1[,1]),
                 f2 = median(js.ms.m1$sims.list$f1[,2]),
                 f3 = median(js.ms.m1$sims.list$f1[,3]),
                 f4 = median(js.ms.m1$sims.list$f1[,4]),
                 f5 = median(js.ms.m1$sims.list$f1[,5]),
                 f6 = median(js.ms.m1$sims.list$f1[,6]),
                 N1 = median(js.ms.m1$sims.list$N[,1]),
                 N2 = median(js.ms.m1$sims.list$N[,2]),
                 N3 = median(js.ms.m1$sims.list$N[,3]),
                 N4 = median(js.ms.m1$sims.list$N[,4]),
                 N5 = median(js.ms.m1$sims.list$N[,5]),
                 N6 = median(js.ms.m1$sims.list$N[,6]),
                 N7 = median(js.ms.m1$sims.list$N[,7]),
                 lambda1 = median(js.ms.m1$sims.list$lambda[,1]),
                 lambda2 = median(js.ms.m1$sims.list$lambda[,2]),
                 lambda3 = median(js.ms.m1$sims.list$lambda[,3]),
                 lambda4 = median(js.ms.m1$sims.list$lambda[,4]),
                 lambda5 = median(js.ms.m1$sims.list$lambda[,5]),
                 lambda6 = median(js.ms.m1$sims.list$lambda[,6])))

    js.sd.df <- rbind(js.sd.df, data.frame(model = "m1",
                 Nsuper = sd(js.ms.m1$sims.list$Nsuper),
                 phi = sd(js.ms.m1$sims.list$mean.phi),
                 b1 = sd(js.ms.m1$sims.list$b[,1]),
                 b2 = sd(js.ms.m1$sims.list$b[,2]),
                 b3 = sd(js.ms.m1$sims.list$b[,3]),
                 b4 = sd(js.ms.m1$sims.list$b[,4]),
                 b5 = sd(js.ms.m1$sims.list$b[,5]),
                 b6 = sd(js.ms.m1$sims.list$b[,6]),
                 b7 = sd(js.ms.m1$sims.list$b[,7]),
                 p = sd(js.ms.m1$sims.list$mean.p),
                 f1 = sd(js.ms.m1$sims.list$f1[,1]),
                 f2 = sd(js.ms.m1$sims.list$f1[,2]),
                 f3 = sd(js.ms.m1$sims.list$f1[,3]),
                 f4 = sd(js.ms.m1$sims.list$f1[,4]),
                 f5 = sd(js.ms.m1$sims.list$f1[,5]),
                 f6 = sd(js.ms.m1$sims.list$f1[,6]),
                 N1 = sd(js.ms.m1$sims.list$N[,1]),
                 N2 = sd(js.ms.m1$sims.list$N[,2]),
                 N3 = sd(js.ms.m1$sims.list$N[,3]),
                 N4 = sd(js.ms.m1$sims.list$N[,4]),
                 N5 = sd(js.ms.m1$sims.list$N[,5]),
                 N6 = sd(js.ms.m1$sims.list$N[,6]),
                 N7 = sd(js.ms.m1$sims.list$N[,7]),
                 lambda1 = sd(js.ms.m1$sims.list$lambda[,1]),
                 lambda2 = sd(js.ms.m1$sims.list$lambda[,2]),
                 lambda3 = sd(js.ms.m1$sims.list$lambda[,3]),
                 lambda4 = sd(js.ms.m1$sims.list$lambda[,4]),
                 lambda5 = sd(js.ms.m1$sims.list$lambda[,5]),
                 lambda6 = sd(js.ms.m1$sims.list$lambda[,6])))

    js.025.df <- rbind(js.025.df, data.frame(model = "m1",
                 Nsuper = quantile(js.ms.m1$sims.list$Nsuper, probs = 0.025),
                 phi = quantile(js.ms.m1$sims.list$mean.phi, probs = 0.025),
                 b1 = quantile(js.ms.m1$sims.list$b[,1], probs = 0.025),
                 b2 = quantile(js.ms.m1$sims.list$b[,2], probs = 0.025),
                 b3 = quantile(js.ms.m1$sims.list$b[,3], probs = 0.025),
                 b4 = quantile(js.ms.m1$sims.list$b[,4], probs = 0.025),
                 b5 = quantile(js.ms.m1$sims.list$b[,5], probs = 0.025),
                 b6 = quantile(js.ms.m1$sims.list$b[,6], probs = 0.025),
                 b7 = quantile(js.ms.m1$sims.list$b[,7], probs = 0.025),
                 p = quantile(js.ms.m1$sims.list$mean.p, probs = 0.025),
                 f1 = quantile(js.ms.m1$sims.list$f1[,1], probs = 0.025),
                 f2 = quantile(js.ms.m1$sims.list$f1[,2], probs = 0.025),
                 f3 = quantile(js.ms.m1$sims.list$f1[,3], probs = 0.025),
                 f4 = quantile(js.ms.m1$sims.list$f1[,4],probs = 0.025),
                 f5 = quantile(js.ms.m1$sims.list$f1[,5], probs = 0.025),
                 f6 = quantile(js.ms.m1$sims.list$f1[,6], probs = 0.025),
                 N1 = quantile(js.ms.m1$sims.list$N[,1], probs = 0.025),
                 N2 = quantile(js.ms.m1$sims.list$N[,2], probs = 0.025),
                 N3 = quantile(js.ms.m1$sims.list$N[,3], probs = 0.025),
                 N4 = quantile(js.ms.m1$sims.list$N[,4], probs = 0.025),
                 N5 = quantile(js.ms.m1$sims.list$N[,5], probs = 0.025),
                 N6 = quantile(js.ms.m1$sims.list$N[,6], probs = 0.025),
                 N7 = quantile(js.ms.m1$sims.list$N[,7], probs = 0.025),
                 lambda1 = quantile(js.ms.m1$sims.list$lambda[,1], probs = 0.025),
                 lambda2 = quantile(js.ms.m1$sims.list$lambda[,2], probs = 0.025),
                 lambda3 = quantile(js.ms.m1$sims.list$lambda[,3], probs = 0.025),
                 lambda4 = quantile(js.ms.m1$sims.list$lambda[,4], probs = 0.025),
                 lambda5 = quantile(js.ms.m1$sims.list$lambda[,5], probs = 0.025),
                 lambda6 = quantile(js.ms.m1$sims.list$lambda[,6], probs = 0.025)))

    js.975.df <- rbind(js.975.df, data.frame(model = "m1",
                 Nsuper = quantile(js.ms.m1$sims.list$Nsuper, probs = 0.975),
                 phi = quantile(js.ms.m1$sims.list$mean.phi, probs = 0.975),
                 b1 = quantile(js.ms.m1$sims.list$b[,1], probs = 0.975),
                 b2 = quantile(js.ms.m1$sims.list$b[,2], probs = 0.975),
                 b3 = quantile(js.ms.m1$sims.list$b[,3], probs = 0.975),
                 b4 = quantile(js.ms.m1$sims.list$b[,4], probs = 0.975),
                 b5 = quantile(js.ms.m1$sims.list$b[,5], probs = 0.975),
                 b6 = quantile(js.ms.m1$sims.list$b[,6], probs = 0.975),
                 b7 = quantile(js.ms.m1$sims.list$b[,7], probs = 0.975),
                 p = quantile(js.ms.m1$sims.list$mean.p, probs = 0.975),
                 f1 = quantile(js.ms.m1$sims.list$f1[,1], probs = 0.975),
                 f2 = quantile(js.ms.m1$sims.list$f1[,2], probs = 0.975),
                 f3 = quantile(js.ms.m1$sims.list$f1[,3], probs = 0.975),
                 f4 = quantile(js.ms.m1$sims.list$f1[,4],probs = 0.975),
                 f5 = quantile(js.ms.m1$sims.list$f1[,5], probs = 0.975),
                 f6 = quantile(js.ms.m1$sims.list$f1[,6], probs = 0.975),
                 N1 = quantile(js.ms.m1$sims.list$N[,1], probs = 0.975),
                 N2 = quantile(js.ms.m1$sims.list$N[,2], probs = 0.975),
                 N3 = quantile(js.ms.m1$sims.list$N[,3], probs = 0.975),
                 N4 = quantile(js.ms.m1$sims.list$N[,4], probs = 0.975),
                 N5 = quantile(js.ms.m1$sims.list$N[,5], probs = 0.975),
                 N6 = quantile(js.ms.m1$sims.list$N[,6], probs = 0.975),
                 N7 = quantile(js.ms.m1$sims.list$N[,7], probs = 0.975),
                 lambda1 = quantile(js.ms.m1$sims.list$lambda[,1], probs = 0.975),
                 lambda2 = quantile(js.ms.m1$sims.list$lambda[,2], probs = 0.975),
                 lambda3 = quantile(js.ms.m1$sims.list$lambda[,3], probs = 0.975),
                 lambda4 = quantile(js.ms.m1$sims.list$lambda[,4], probs = 0.975),
                 lambda5 = quantile(js.ms.m1$sims.list$lambda[,5], probs = 0.975),
                 lambda6 = quantile(js.ms.m1$sims.list$lambda[,6], probs = 0.975)))

    js.mcmc.df <- rbind(js.mcmc.df, data.frame(model="m1",
                  rt=js.ms.m1$mcmc.info$elapsed.mins,
                  min.ess=min(unlist(js.ms.m1$n.eff)),
                  max.ess=max(unlist(js.ms.m1$n.eff)),
                  med.ess=median(unlist(js.ms.m1$n.eff)),
                  max.Rhat=max(unlist(js.ms.m1$Rhat))))


    # model m2 -------------------------------------------------------

    all_add<-function(NO_OCC=7){
      # calculates different combinations of arrival and departure timing
      arr<-numeric()
      dep<-numeric()
      t<-1
      for (i in 1:NO_OCC){
        for (j in i:NO_OCC){
          arr[t]<-i
          dep[t]<-j
          t<-t+1
        }}
      return(cbind(arr,dep))}

    MS=dim(ys)[1]

    poss<-all_add()

    first<-function(x){
      #determine timing of first observation
      which(x==1)[1]}

    last<-function(x){
      #determine timing of last observation
      tx<-which(x==1)
      lx<-length(tx)
      tx[lx]
    }

    fd<-apply(ys[-MS,-1],1,first)

    ld<-apply(ys[-MS,-1],1,last)

    nposs=length(poss[,1])

    poss_obs<-matrix(NA,nrow=nposs,ncol=(MS-1))

    for (i in 1:(MS-1)){
      #for each summarized capture history this loop determines if the possible
      # underlying dynamics implied by poss are possible
      poss_obs[,i]<-ifelse(poss[,1]<=fd[i]&poss[,2]>=ld[i],1,0)
    }

    Nobs_ind <- sum(FR[-MS])

    lookup <- rep(seq(1,length(FR[-MS]), 1), FR[-MS])

    jags.data<- list(FR=FR, fr=FR, ys = ys, n.occasions=dim(ys)[2], MS=dim(ys)[1],
                     Nobs_ind = Nobs_ind,Npos_ind = sum(FR) , lookup=lookup,
                     nposs=nposs, poss=poss, poss_obs=poss_obs)

    inits <- function(){list(mean.phi = runif(1, .6, .8),
                             mean.p = runif(1, .4, .6), A=rep(c(fd,NA),FR),
                             L=rep(c(ld,NA),FR), ar=rep(c(fd,1), FR),
                             li=rep(c(ld,(dim(ys)[2]-1)),FR), U=FR[MS])}

    js.ms.m2 <- jags(jags.data, inits, parameters, "js-ms-m2.txt", n.chains = nc,
                     n.thin =nt, n.iter = ni, n.burnin = nb, parallel=TRUE)

    # extract results
    js.med.df <- rbind(js.med.df, data.frame(model = "m2",
                 Nsuper = median(js.ms.m2$sims.list$Nsuper),
                 phi = median(js.ms.m2$sims.list$mean.phi),
                 b1 = median(js.ms.m2$sims.list$b[,1]),
                 b2 = median(js.ms.m2$sims.list$b[,2]),
                 b3 = median(js.ms.m2$sims.list$b[,3]),
                 b4 = median(js.ms.m2$sims.list$b[,4]),
                 b5 = median(js.ms.m2$sims.list$b[,5]),
                 b6 = median(js.ms.m2$sims.list$b[,6]),
                 b7 = median(js.ms.m2$sims.list$b[,7]),
                 p = median(js.ms.m2$sims.list$mean.p),
                 f1 = median(js.ms.m2$sims.list$f1[,1]),
                 f2 = median(js.ms.m2$sims.list$f1[,2]),
                 f3 = median(js.ms.m2$sims.list$f1[,3]),
                 f4 = median(js.ms.m2$sims.list$f1[,4]),
                 f5 = median(js.ms.m2$sims.list$f1[,5]),
                 f6 = median(js.ms.m2$sims.list$f1[,6]),
                 N1 = median(js.ms.m2$sims.list$N[,1]),
                 N2 = median(js.ms.m2$sims.list$N[,2]),
                 N3 = median(js.ms.m2$sims.list$N[,3]),
                 N4 = median(js.ms.m2$sims.list$N[,4]),
                 N5 = median(js.ms.m2$sims.list$N[,5]),
                 N6 = median(js.ms.m2$sims.list$N[,6]),
                 N7 = median(js.ms.m2$sims.list$N[,7]),
                 lambda1 = median(js.ms.m2$sims.list$lambda[,1]),
                 lambda2 = median(js.ms.m2$sims.list$lambda[,2]),
                 lambda3 = median(js.ms.m2$sims.list$lambda[,3]),
                 lambda4 = median(js.ms.m2$sims.list$lambda[,4]),
                 lambda5 = median(js.ms.m2$sims.list$lambda[,5]),
                 lambda6 = median(js.ms.m2$sims.list$lambda[,6])))

    js.sd.df <- rbind(js.sd.df, data.frame(model = "m2",
                Nsuper = sd(js.ms.m2$sims.list$Nsuper),
                phi = sd(js.ms.m2$sims.list$mean.phi),
                b1 = sd(js.ms.m2$sims.list$b[,1]),
                b2 = sd(js.ms.m2$sims.list$b[,2]),
                b3 = sd(js.ms.m2$sims.list$b[,3]),
                b4 = sd(js.ms.m2$sims.list$b[,4]),
                b5 = sd(js.ms.m2$sims.list$b[,5]),
                b6 = sd(js.ms.m2$sims.list$b[,6]),
                b7 = sd(js.ms.m2$sims.list$b[,7]),
                p = sd(js.ms.m2$sims.list$mean.p),
                f1 = sd(js.ms.m2$sims.list$f1[,1]),
                f2 = sd(js.ms.m2$sims.list$f1[,2]),
                f3 = sd(js.ms.m2$sims.list$f1[,3]),
                f4 = sd(js.ms.m2$sims.list$f1[,4]),
                f5 = sd(js.ms.m2$sims.list$f1[,5]),
                f6 = sd(js.ms.m2$sims.list$f1[,6]),
                N1 = sd(js.ms.m2$sims.list$N[,1]),
                N2 = sd(js.ms.m2$sims.list$N[,2]),
                N3 = sd(js.ms.m2$sims.list$N[,3]),
                N4 = sd(js.ms.m2$sims.list$N[,4]),
                N5 = sd(js.ms.m2$sims.list$N[,5]),
                N6 = sd(js.ms.m2$sims.list$N[,6]),
                N7 = sd(js.ms.m2$sims.list$N[,7]),
                lambda1 = sd(js.ms.m2$sims.list$lambda[,1]),
                lambda2 = sd(js.ms.m2$sims.list$lambda[,2]),
                lambda3 = sd(js.ms.m2$sims.list$lambda[,3]),
                lambda4 = sd(js.ms.m2$sims.list$lambda[,4]),
                lambda5 = sd(js.ms.m2$sims.list$lambda[,5]),
                lambda6 = sd(js.ms.m2$sims.list$lambda[,6])))

    js.025.df <- rbind(js.025.df, data.frame(model = "m2",
                 Nsuper = quantile(js.ms.m2$sims.list$Nsuper, probs = 0.025),
                 phi = quantile(js.ms.m2$sims.list$mean.phi, probs = 0.025),
                 b1 = quantile(js.ms.m2$sims.list$b[,1], probs = 0.025),
                 b2 = quantile(js.ms.m2$sims.list$b[,2], probs = 0.025),
                 b3 = quantile(js.ms.m2$sims.list$b[,3], probs = 0.025),
                 b4 = quantile(js.ms.m2$sims.list$b[,4], probs = 0.025),
                 b5 = quantile(js.ms.m2$sims.list$b[,5], probs = 0.025),
                 b6 = quantile(js.ms.m2$sims.list$b[,6], probs = 0.025),
                 b7 = quantile(js.ms.m2$sims.list$b[,7], probs = 0.025),
                 p = quantile(js.ms.m2$sims.list$mean.p, probs = 0.025),
                 f1 = quantile(js.ms.m2$sims.list$f1[,1], probs = 0.025),
                 f2 = quantile(js.ms.m2$sims.list$f1[,2], probs = 0.025),
                 f3 = quantile(js.ms.m2$sims.list$f1[,3], probs = 0.025),
                 f4 = quantile(js.ms.m2$sims.list$f1[,4], probs = 0.025),
                 f5 = quantile(js.ms.m2$sims.list$f1[,5], probs = 0.025),
                 f6 = quantile(js.ms.m2$sims.list$f1[,6], probs = 0.025),
                 N1 = quantile(js.ms.m2$sims.list$N[,1], probs = 0.025),
                 N2 = quantile(js.ms.m2$sims.list$N[,2], probs = 0.025),
                 N3 = quantile(js.ms.m2$sims.list$N[,3], probs = 0.025),
                 N4 = quantile(js.ms.m2$sims.list$N[,4], probs = 0.025),
                 N5 = quantile(js.ms.m2$sims.list$N[,5], probs = 0.025),
                 N6 = quantile(js.ms.m2$sims.list$N[,6], probs = 0.025),
                 N7 = quantile(js.ms.m2$sims.list$N[,7], probs = 0.025),
                 lambda1 = quantile(js.ms.m2$sims.list$lambda[,1], probs = 0.025),
                 lambda2 = quantile(js.ms.m2$sims.list$lambda[,2], probs = 0.025),
                 lambda3 = quantile(js.ms.m2$sims.list$lambda[,3], probs = 0.025),
                 lambda4 = quantile(js.ms.m2$sims.list$lambda[,4], probs = 0.025),
                 lambda5 = quantile(js.ms.m2$sims.list$lambda[,5], probs = 0.025),
                 lambda6 = quantile(js.ms.m2$sims.list$lambda[,6], probs = 0.025)))

    js.975.df <- rbind(js.975.df, data.frame(model = "m2",
                 Nsuper = quantile(js.ms.m2$sims.list$Nsuper, probs = 0.975),
                 phi = quantile(js.ms.m2$sims.list$mean.phi, probs = 0.975),
                 b1 = quantile(js.ms.m2$sims.list$b[,1], probs = 0.975),
                 b2 = quantile(js.ms.m2$sims.list$b[,2], probs = 0.975),
                 b3 = quantile(js.ms.m2$sims.list$b[,3], probs = 0.975),
                 b4 = quantile(js.ms.m2$sims.list$b[,4], probs = 0.975),
                 b5 = quantile(js.ms.m2$sims.list$b[,5], probs = 0.975),
                 b6 = quantile(js.ms.m2$sims.list$b[,6], probs = 0.975),
                 b7 = quantile(js.ms.m2$sims.list$b[,7], probs = 0.975),
                 p = quantile(js.ms.m2$sims.list$mean.p, probs = 0.975),
                 f1 = quantile(js.ms.m2$sims.list$f1[,1], probs = 0.975),
                 f2 = quantile(js.ms.m2$sims.list$f1[,2], probs = 0.975),
                 f3 = quantile(js.ms.m2$sims.list$f1[,3], probs = 0.975),
                 f4 = quantile(js.ms.m2$sims.list$f1[,4], probs = 0.975),
                 f5 = quantile(js.ms.m2$sims.list$f1[,5], probs = 0.975),
                 f6 = quantile(js.ms.m2$sims.list$f1[,6], probs = 0.975),
                 N1 = quantile(js.ms.m2$sims.list$N[,1], probs = 0.975),
                 N2 = quantile(js.ms.m2$sims.list$N[,2], probs = 0.975),
                 N3 = quantile(js.ms.m2$sims.list$N[,3], probs = 0.975),
                 N4 = quantile(js.ms.m2$sims.list$N[,4], probs = 0.975),
                 N5 = quantile(js.ms.m2$sims.list$N[,5], probs = 0.975),
                 N6 = quantile(js.ms.m2$sims.list$N[,6], probs = 0.975),
                 N7 = quantile(js.ms.m2$sims.list$N[,7], probs = 0.975),
                 lambda1 = quantile(js.ms.m2$sims.list$lambda[,1], probs = 0.975),
                 lambda2 = quantile(js.ms.m2$sims.list$lambda[,2], probs = 0.975),
                 lambda3 = quantile(js.ms.m2$sims.list$lambda[,3], probs = 0.975),
                 lambda4 = quantile(js.ms.m2$sims.list$lambda[,4], probs = 0.975),
                 lambda5 = quantile(js.ms.m2$sims.list$lambda[,5], probs = 0.975),
                 lambda6 = quantile(js.ms.m2$sims.list$lambda[,6], probs = 0.975)))

    js.mcmc.df <- rbind(js.mcmc.df, data.frame(model="m2",
                  rt=js.ms.m2$mcmc.info$elapsed.mins,
                  min.ess=min(unlist(js.ms.m2$n.eff)),
                  max.ess=max(unlist(js.ms.m2$n.eff)),
                  med.ess=median(unlist(js.ms.m2$n.eff)),
                  max.Rhat=max(unlist(js.ms.m2$Rhat))))


    # model m3 -------------------------------------------------------

    # create index to recover discrete latent states for derived parameters
    lookup <- rep(seq(1,length(FR), 1), FR)
    jags.data<- list(FR=FR, fr=FR, ys = ys, n.occasions=dim(ys)[2], MS=dim(ys)[1], M=M,
                     lookup=lookup)
    inits <- function(){list(mean.phi = runif(1, .6, .8), mean.p = runif(1, .4, .6),
                             U=FR[MS])}
    js.ms.m3 <- jags(jags.data, inits, parameters, "js-ms-m3.txt", n.chains = nc,
                     n.thin =nt, n.iter = ni, n.burnin = nb, parallel=TRUE)

    # extract results
    js.med.df <- rbind(js.med.df, data.frame(model = "m3",
                 Nsuper = median(js.ms.m3$sims.list$Nsuper),
                 phi = median(js.ms.m3$sims.list$mean.phi),
                 b1 = median(js.ms.m3$sims.list$b[,1]),
                 b2 = median(js.ms.m3$sims.list$b[,2]),
                 b3 = median(js.ms.m3$sims.list$b[,3]),
                 b4 = median(js.ms.m3$sims.list$b[,4]),
                 b5 = median(js.ms.m3$sims.list$b[,5]),
                 b6 = median(js.ms.m3$sims.list$b[,6]),
                 b7 = median(js.ms.m3$sims.list$b[,7]),
                 p = median(js.ms.m3$sims.list$mean.p),
                 f1 = median(js.ms.m3$sims.list$f1[,1]),
                 f2 = median(js.ms.m3$sims.list$f1[,2]),
                 f3 = median(js.ms.m3$sims.list$f1[,3]),
                 f4 = median(js.ms.m3$sims.list$f1[,4]),
                 f5 = median(js.ms.m3$sims.list$f1[,5]),
                 f6 = median(js.ms.m3$sims.list$f1[,6]),
                 N1 = median(js.ms.m3$sims.list$N[,1]),
                 N2 = median(js.ms.m3$sims.list$N[,2]),
                 N3 = median(js.ms.m3$sims.list$N[,3]),
                 N4 = median(js.ms.m3$sims.list$N[,4]),
                 N5 = median(js.ms.m3$sims.list$N[,5]),
                 N6 = median(js.ms.m3$sims.list$N[,6]),
                 N7 = median(js.ms.m3$sims.list$N[,7]),
                 lambda1 = median(js.ms.m3$sims.list$lambda[,1]),
                 lambda2 = median(js.ms.m3$sims.list$lambda[,2]),
                 lambda3 = median(js.ms.m3$sims.list$lambda[,3]),
                 lambda4 = median(js.ms.m3$sims.list$lambda[,4]),
                 lambda5 = median(js.ms.m3$sims.list$lambda[,5]),
                 lambda6 = median(js.ms.m3$sims.list$lambda[,6])))

    js.sd.df <- rbind(js.sd.df, data.frame(model = "m3",
                Nsuper = sd(js.ms.m3$sims.list$Nsuper),
                phi = sd(js.ms.m3$sims.list$mean.phi),
                b1 = sd(js.ms.m3$sims.list$b[,1]),
                b2 = sd(js.ms.m3$sims.list$b[,2]),
                b3 = sd(js.ms.m3$sims.list$b[,3]),
                b4 = sd(js.ms.m3$sims.list$b[,4]),
                b5 = sd(js.ms.m3$sims.list$b[,5]),
                b6 = sd(js.ms.m3$sims.list$b[,6]),
                b7 = sd(js.ms.m3$sims.list$b[,7]),
                p = sd(js.ms.m3$sims.list$mean.p),
                f1 = sd(js.ms.m3$sims.list$f1[,1]),
                f2 = sd(js.ms.m3$sims.list$f1[,2]),
                f3 = sd(js.ms.m3$sims.list$f1[,3]),
                f4 = sd(js.ms.m3$sims.list$f1[,4]),
                f5 = sd(js.ms.m3$sims.list$f1[,5]),
                f6 = sd(js.ms.m3$sims.list$f1[,6]),
                N1 = sd(js.ms.m3$sims.list$N[,1]),
                N2 = sd(js.ms.m3$sims.list$N[,2]),
                N3 = sd(js.ms.m3$sims.list$N[,3]),
                N4 = sd(js.ms.m3$sims.list$N[,4]),
                N5 = sd(js.ms.m3$sims.list$N[,5]),
                N6 = sd(js.ms.m3$sims.list$N[,6]),
                N7 = sd(js.ms.m3$sims.list$N[,7]),
                lambda1 = sd(js.ms.m3$sims.list$lambda[,1]),
                lambda2 = sd(js.ms.m3$sims.list$lambda[,2]),
                lambda3 = sd(js.ms.m3$sims.list$lambda[,3]),
                lambda4 = sd(js.ms.m3$sims.list$lambda[,4]),
                lambda5 = sd(js.ms.m3$sims.list$lambda[,5]),
                lambda6 = sd(js.ms.m3$sims.list$lambda[,6])))

    js.025.df <- rbind(js.025.df, data.frame(model = "m3",
                 Nsuper = quantile(js.ms.m3$sims.list$Nsuper, probs = 0.025),
                 phi = quantile(js.ms.m3$sims.list$mean.phi, probs = 0.025),
                 b1 = quantile(js.ms.m3$sims.list$b[,1], probs = 0.025),
                 b2 = quantile(js.ms.m3$sims.list$b[,2], probs = 0.025),
                 b3 = quantile(js.ms.m3$sims.list$b[,3], probs = 0.025),
                 b4 = quantile(js.ms.m3$sims.list$b[,4], probs = 0.025),
                 b5 = quantile(js.ms.m3$sims.list$b[,5], probs = 0.025),
                 b6 = quantile(js.ms.m3$sims.list$b[,6], probs = 0.025),
                 b7 = quantile(js.ms.m3$sims.list$b[,7], probs = 0.025),
                 p = quantile(js.ms.m3$sims.list$mean.p, probs = 0.025),
                 f1 = quantile(js.ms.m3$sims.list$f1[,1], probs = 0.025),
                 f2 = quantile(js.ms.m3$sims.list$f1[,2], probs = 0.025),
                 f3 = quantile(js.ms.m3$sims.list$f1[,3], probs = 0.025),
                 f4 = quantile(js.ms.m3$sims.list$f1[,4], probs = 0.025),
                 f5 = quantile(js.ms.m3$sims.list$f1[,5], probs = 0.025),
                 f6 = quantile(js.ms.m3$sims.list$f1[,6], probs = 0.025),
                 N1 = quantile(js.ms.m3$sims.list$N[,1], probs = 0.025),
                 N2 = quantile(js.ms.m3$sims.list$N[,2], probs = 0.025),
                 N3 = quantile(js.ms.m3$sims.list$N[,3], probs = 0.025),
                 N4 = quantile(js.ms.m3$sims.list$N[,4], probs = 0.025),
                 N5 = quantile(js.ms.m3$sims.list$N[,5], probs = 0.025),
                 N6 = quantile(js.ms.m3$sims.list$N[,6], probs = 0.025),
                 N7 = quantile(js.ms.m3$sims.list$N[,7], probs = 0.025),
                 lambda1 = quantile(js.ms.m3$sims.list$lambda[,1], probs = 0.025),
                 lambda2 = quantile(js.ms.m3$sims.list$lambda[,2], probs = 0.025),
                 lambda3 = quantile(js.ms.m3$sims.list$lambda[,3], probs = 0.025),
                 lambda4 = quantile(js.ms.m3$sims.list$lambda[,4], probs = 0.025),
                 lambda5 = quantile(js.ms.m3$sims.list$lambda[,5], probs = 0.025),
                 lambda6 = quantile(js.ms.m3$sims.list$lambda[,6], probs = 0.025)))

    js.975.df <- rbind(js.975.df, data.frame(model = "m3",
                 Nsuper = quantile(js.ms.m3$sims.list$Nsuper, probs = 0.975),
                 phi = quantile(js.ms.m3$sims.list$mean.phi, probs = 0.975),
                 b1 = quantile(js.ms.m3$sims.list$b[,1], probs = 0.975),
                 b2 = quantile(js.ms.m3$sims.list$b[,2], probs = 0.975),
                 b3 = quantile(js.ms.m3$sims.list$b[,3], probs = 0.975),
                 b4 = quantile(js.ms.m3$sims.list$b[,4], probs = 0.975),
                 b5 = quantile(js.ms.m3$sims.list$b[,5], probs = 0.975),
                 b6 = quantile(js.ms.m3$sims.list$b[,6], probs = 0.975),
                 b7 = quantile(js.ms.m3$sims.list$b[,7], probs = 0.975),
                 p = quantile(js.ms.m3$sims.list$mean.p, probs = 0.975),
                 f1 = quantile(js.ms.m3$sims.list$f1[,1], probs = 0.975),
                 f2 = quantile(js.ms.m3$sims.list$f1[,2], probs = 0.975),
                 f3 = quantile(js.ms.m3$sims.list$f1[,3], probs = 0.975),
                 f4 = quantile(js.ms.m3$sims.list$f1[,4], probs = 0.975),
                 f5 = quantile(js.ms.m3$sims.list$f1[,5], probs = 0.975),
                 f6 = quantile(js.ms.m3$sims.list$f1[,6], probs = 0.975),
                 N1 = quantile(js.ms.m3$sims.list$N[,1], probs = 0.975),
                 N2 = quantile(js.ms.m3$sims.list$N[,2], probs = 0.975),
                 N3 = quantile(js.ms.m3$sims.list$N[,3], probs = 0.975),
                 N4 = quantile(js.ms.m3$sims.list$N[,4], probs = 0.975),
                 N5 = quantile(js.ms.m3$sims.list$N[,5], probs = 0.975),
                 N6 = quantile(js.ms.m3$sims.list$N[,6], probs = 0.975),
                 N7 = quantile(js.ms.m3$sims.list$N[,7], probs = 0.975),
                 lambda1 = quantile(js.ms.m3$sims.list$lambda[,1], probs = 0.975),
                 lambda2 = quantile(js.ms.m3$sims.list$lambda[,2], probs = 0.975),
                 lambda3 = quantile(js.ms.m3$sims.list$lambda[,3], probs = 0.975),
                 lambda4 = quantile(js.ms.m3$sims.list$lambda[,4], probs = 0.975),
                 lambda5 = quantile(js.ms.m3$sims.list$lambda[,5], probs = 0.975),
                 lambda6 = quantile(js.ms.m3$sims.list$lambda[,6], probs = 0.975)))

    js.mcmc.df <- rbind(js.mcmc.df, data.frame(model="m3",
                  rt=js.ms.m3$mcmc.info$elapsed.mins,
                  min.ess=min(unlist(js.ms.m3$n.eff)),
                  max.ess=max(unlist(js.ms.m3$n.eff)),
                  med.ess=median(unlist(js.ms.m3$n.eff)),
                  max.Rhat=max(unlist(js.ms.m3$Rhat))))
  }
  return(list(js.med.df, js.sd.df, js.025.df, js.975.df, js.mcmc.df))
}

if (file.exists("js_out_marg.rds")) {
  js.out.marg <- readRDS("js_out_marg.rds")
} else {
  js.out.marg <- js(nrep)  # a time-consuming function
  saveRDS(js.out.marg, "js_out_marg.rds")
}

## extract posterior medians
d.meds <- js.out.marg[[1]][js.out.marg[[1]]$model%in%"d",]
m1.meds <- js.out.marg[[1]][js.out.marg[[1]]$model%in%"m1",]
m2.meds <- js.out.marg[[1]][js.out.marg[[1]]$model%in%"m2",]
m3.meds <- js.out.marg[[1]][js.out.marg[[1]]$model%in%"m3",]

save(d.meds, m1.meds, m2.meds, m3.meds, file = "modelcomparefigS1.RData")

library(kableExtra)
options(knitr.kable.NA = '')

js.out.marg[[5]]$mps100 <- js.out.marg[[5]]$rt/js.out.marg[[5]]$min.ess*100
js.out.marg[[5]]$mps1000 <- js.out.marg[[5]]$rt/js.out.marg[[5]]$min.ess*1000

# expected values
E_N <- c(136, 139, 141, 143, 144, 145, 145)
E_lam <- E_N[2:7]/E_N[1:6]
E_f <- E_lam - .7

# summarize simulations
simsum <- data.frame(model = js.out.marg[[3]]$model,
                     Nsuper.cov = ifelse(js.out.marg[[3]]$Nsuper <= 400 &
                                           js.out.marg[[4]]$Nsuper >= 400, 1, 0),
                     Nsuper.w = js.out.marg[[4]]$Nsuper - js.out.marg[[3]]$Nsuper,
                     N1.cov = ifelse(js.out.marg[[3]]$N1 <= E_N[1] &
                                       js.out.marg[[4]]$N1 >= E_N[1], 1, 0),
                     N1.w = js.out.marg[[4]]$N1 - js.out.marg[[3]]$N1,
                     N2.cov = ifelse(js.out.marg[[3]]$N2 <= E_N[2] &
                                       js.out.marg[[4]]$N2 >= E_N[2], 1, 0),
                     N2.w = js.out.marg[[4]]$N2 - js.out.marg[[3]]$N2,
                     N3.cov = ifelse(js.out.marg[[3]]$N3 <= E_N[3] &
                                       js.out.marg[[4]]$N3 >= E_N[3], 1, 0),
                     N3.w = js.out.marg[[4]]$N3 - js.out.marg[[3]]$N3,
                     N4.cov = ifelse(js.out.marg[[3]]$N4 <= E_N[4] &
                                       js.out.marg[[4]]$N4 >= E_N[4], 1, 0),
                     N4.w = js.out.marg[[4]]$N4 - js.out.marg[[3]]$N4,
                     N5.cov = ifelse(js.out.marg[[3]]$N5 <= E_N[5] &
                                       js.out.marg[[4]]$N5 >= E_N[5], 1, 0),
                     N5.w = js.out.marg[[4]]$N5 - js.out.marg[[3]]$N5,
                     N6.cov = ifelse(js.out.marg[[3]]$N6 <= E_N[6] &
                                       js.out.marg[[4]]$N6 >= E_N[6], 1, 0),
                     N6.w = js.out.marg[[4]]$N6 - js.out.marg[[3]]$N6,
                     N7.cov = ifelse(js.out.marg[[3]]$N7 <= E_N[7] &
                                       js.out.marg[[4]]$N7 >= E_N[7], 1, 0),
                     N7.w = js.out.marg[[4]]$N7 - js.out.marg[[3]]$N7,
                     lam1.cov = ifelse(js.out.marg[[3]]$lambda1 <= E_lam[1] &
                                         js.out.marg[[4]]$lambda1 >= E_lam[1], 1, 0),
                     lam1.w = js.out.marg[[4]]$lambda1 - js.out.marg[[3]]$lambda1,
                     lam2.cov = ifelse(js.out.marg[[3]]$lambda2 <= E_lam[2] &
                                         js.out.marg[[4]]$lambda2 >= E_lam[2], 1, 0),
                     lam2.w = js.out.marg[[4]]$lambda2 - js.out.marg[[3]]$lambda2,
                     lam3.cov = ifelse(js.out.marg[[3]]$lambda3 <= E_lam[3] &
                                         js.out.marg[[4]]$lambda3 >= E_lam[3], 1, 0),
                     lam3.w = js.out.marg[[4]]$lambda3 - js.out.marg[[3]]$lambda3,
                     lam4.cov = ifelse(js.out.marg[[3]]$lambda4 <= E_lam[4] &
                                         js.out.marg[[4]]$lambda4 >= E_lam[4], 1, 0),
                     lam4.w = js.out.marg[[4]]$lambda4 - js.out.marg[[3]]$lambda4,
                     lam5.cov = ifelse(js.out.marg[[3]]$lambda5 <= E_lam[5] &
                                         js.out.marg[[4]]$lambda5 >= E_lam[5], 1, 0),
                     lam5.w = js.out.marg[[4]]$lambda5 - js.out.marg[[3]]$lambda5,
                     lam6.cov = ifelse(js.out.marg[[3]]$lambda6 <= E_lam[6] &
                                         js.out.marg[[4]]$lambda6 >= E_lam[6], 1, 0),
                     lam6.w = js.out.marg[[4]]$lambda6 - js.out.marg[[3]]$lambda6,
                     f1.cov = ifelse(js.out.marg[[3]]$f1 <= E_f[1] &
                                       js.out.marg[[4]]$f1 >= E_f[1], 1, 0),
                     f1.w = js.out.marg[[4]]$f1 - js.out.marg[[3]]$f1,
                     f2.cov = ifelse(js.out.marg[[3]]$f2 <= E_f[2] &
                                       js.out.marg[[4]]$f2 >= E_f[2], 1, 0),
                     f2.w = js.out.marg[[4]]$f2 - js.out.marg[[3]]$f2,
                     f3.cov = ifelse(js.out.marg[[3]]$f3 <= E_f[3] &
                                       js.out.marg[[4]]$f3 >= E_f[3], 1, 0),
                     f3.w = js.out.marg[[4]]$f3 - js.out.marg[[3]]$f3,
                     f4.cov = ifelse(js.out.marg[[3]]$f4 <= E_f[4] &
                                       js.out.marg[[4]]$f4 >= E_f[4], 1, 0),
                     f4.w = js.out.marg[[4]]$f4 - js.out.marg[[3]]$f4,
                     f5.cov = ifelse(js.out.marg[[3]]$f5 <= E_f[5] &
                                       js.out.marg[[4]]$f5 >= E_f[5], 1, 0),
                     f5.w = js.out.marg[[4]]$f5 - js.out.marg[[3]]$f5,
                     f6.cov = ifelse(js.out.marg[[3]]$f6 <= E_f[6] &
                                       js.out.marg[[4]]$f6 >= E_f[6], 1, 0),
                     f6.w = js.out.marg[[4]]$f6 - js.out.marg[[3]]$f6)

library(tidyverse)
library(data.table)

x <- simsum %>% group_by(model) %>% summarise(across(.cols=everything(),
                          list(mean = mean, sd = sd), .names = "{.col}.{.fn}"))

xt <- data.table::transpose(x, make.names="model")
xt$parm <- unlist(strsplit(names(x)[2:ncol(x)], "\\.", ""))[seq(1, length(unlist(strsplit(names(x)[2:ncol(x)], "\\.", ""))), 3)]
xt$var <- rep(c("cov", "cov", "w", "w"), 20)
xt$stat <- rep(c("mean", "sd"), 40)
xt <- xt %>% pivot_wider(id_cols = c(parm, stat), names_from = var, values_from = c(d, m1, m2, m3))
xt <- data.frame(xt)

xt$d_cov <- sprintf("%.1f", round(xt$d_cov*100, 1))
xt$d_w[xt$parm %in% c("Nsuper", "N1", "N2", "N3", "N4", "N5", "N6", "N7")] <-
  sprintf("%.1f", round(xt$d_w[xt$parm %in% c("Nsuper", "N1", "N2", "N3", "N4",
                                              "N5", "N6", "N7")], 1))
xt$d_w[!(xt$parm %in% c("Nsuper", "N1", "N2", "N3", "N4", "N5", "N6", "N7"))] <-
  sprintf("%.3f", round(as.numeric(xt$d_w[!(xt$parm %in% c("Nsuper", "N1",
                                    "N2", "N3", "N4", "N5", "N6", "N7"))]), 3))

xt$m1_cov <- sprintf("%.1f", round(xt$m1_cov*100, 1))
xt$m1_w[xt$parm %in% c("Nsuper", "N1", "N2", "N3", "N4", "N5", "N6", "N7")] <-
  sprintf("%.1f", round(xt$m1_w[xt$parm %in% c("Nsuper", "N1", "N2", "N3", "N4",
                                               "N5", "N6", "N7")], 1))
xt$m1_w[!(xt$parm %in% c("Nsuper", "N1", "N2", "N3", "N4", "N5", "N6", "N7"))] <-
  sprintf("%.3f", round(as.numeric(xt$m1_w[!(xt$parm %in% c("Nsuper", "N1", "N2",
                                          "N3", "N4", "N5", "N6", "N7"))]), 3))

xt$m2_cov <- sprintf("%.1f", round(xt$m2_cov*100, 1))
xt$m2_w[xt$parm %in% c("Nsuper", "N1", "N2", "N3", "N4", "N5", "N6", "N7")] <-
  sprintf("%.1f", round(xt$m2_w[xt$parm %in% c("Nsuper", "N1", "N2", "N3", "N4",
                                               "N5", "N6", "N7")], 1))
xt$m2_w[!(xt$parm %in% c("Nsuper", "N1", "N2", "N3", "N4", "N5", "N6", "N7"))] <-
  sprintf("%.3f", round(as.numeric(xt$m2_w[!(xt$parm %in% c("Nsuper", "N1", "N2",
                                          "N3", "N4", "N5", "N6", "N7"))]), 3))

xt$m3_cov <- sprintf("%.1f", round(xt$m3_cov*100, 1))
xt$m3_w[xt$parm %in% c("Nsuper", "N1", "N2", "N3", "N4", "N5", "N6", "N7")] <-
  sprintf("%.1f", round(xt$m3_w[xt$parm %in% c("Nsuper", "N1", "N2", "N3", "N4",
                                               "N5", "N6", "N7")], 1))
xt$m3_w[!(xt$parm %in% c("Nsuper", "N1", "N2", "N3", "N4", "N5", "N6", "N7"))] <-
  sprintf("%.3f", round(as.numeric(xt$m3_w[!(xt$parm %in% c("Nsuper", "N1", "N2",
                                          "N3", "N4", "N5", "N6", "N7"))]), 3))

xt$parm[seq(2,40,2)] <- NA

xt$d_cov[seq(2,40,2)] <- paste0("(", xt$d_cov[seq(2,40,2)], ")")
xt$d_w[seq(2,40,2)] <- paste0("(", xt$d_w[seq(2,40,2)], ")")
xt$m1_cov[seq(2,40,2)] <- paste0("(", xt$m1_cov[seq(2,40,2)], ")")
xt$m1_w[seq(2,40,2)] <- paste0("(", xt$m1_w[seq(2,40,2)], ")")
xt$m2_cov[seq(2,40,2)] <- paste0("(", xt$m2_cov[seq(2,40,2)], ")")
xt$m2_w[seq(2,40,2)] <- paste0("(", xt$m2_w[seq(2,40,2)], ")")
xt$m3_cov[seq(2,40,2)] <- paste0("(", xt$m3_cov[seq(2,40,2)], ")")
xt$m3_w[seq(2,40,2)] <- paste0("(", xt$m3_w[seq(2,40,2)], ")")

tabS1 <- data.frame(xt[,c(1,3:10)])
tabS1$parm[1] <- "$N_{super}$"
tabS1$parm[3] <- "$N_{1}$"
tabS1$parm[5] <- "$N_{2}$"
tabS1$parm[7] <- "$N_{3}$"
tabS1$parm[9] <- "$N_{4}$"
tabS1$parm[11] <- "$N_{5}$"
tabS1$parm[13] <- "$N_{6}$"
tabS1$parm[15] <- "$N_{7}$"
tabS1$parm[17] <- "$\\lambda_1$"
tabS1$parm[19] <- "$\\lambda_2$"
tabS1$parm[21] <- "$\\lambda_3$"
tabS1$parm[23] <- "$\\lambda_4$"
tabS1$parm[25] <- "$\\lambda_5$"
tabS1$parm[27] <- "$\\lambda_6$"
tabS1$parm[29] <- "$f_1$"
tabS1$parm[31] <- "$f_2$"
tabS1$parm[33] <- "$f_3$"
tabS1$parm[35] <- "$f_4$"
tabS1$parm[37] <- "$f_5$"
tabS1$parm[39] <- "$f_6$"

```

#-------------------------------------------------------------------------------------------------------------------
\# Create Table S1

kbl(tabS1, format = "pipe", booktabs = TRUE, caption = "Table S1. 95\\% credible interval coverage and widths [mean (sd)] for discrete (d) and marginalized (m1, m2, and m3) JS models applied to 100 simulated data sets that differed with respect to how abundance (N), population change, and recruitment are estimated.", col.names = c("", "\\% coverage", "Width", "\\% coverage", "Width", "\\% coverage", "Width", "\\% coverage", "Width"), align = "c") %>%
 add_header_above(header = c(" ", "d" = 2, "m1" = 2, "m2" = 2, "m3" = 2))

\# note that “add_header_above() not working when knitting document to Word.

#----------------------------------------------------------------------------

\# Create Figure S1

```
library(fields)
library(wesanderson)

par(family = "serif", mfrow=c(3,1), mar = c(4,5,0,3))
# in response to review, ad hoc y-axis split, Nsuper shown on right
d.meds$Nsuper <- d.meds$Nsuper - 100
m1.meds$Nsuper <- m1.meds$Nsuper - 100
m2.meds$Nsuper <- m2.meds$Nsuper - 100
m3.meds$Nsuper <- m3.meds$Nsuper - 100

bplot(d.meds[,c(18:24,2)], use.cols = TRUE, outline = FALSE, staplewex=0, col= wes_palette("Royal1")[1],
      boxwex=.15, lty=1, pos = seq(from=.5, by=2, length=8), xaxt="n", yaxt="n", ylab = "Abundance (N)",
      xlim=c(0,16), ylim=c(50, 400), las=1, cex.lab=1.25, cex.axis=1)
axis(2, at = c(100, 150, 200), las=1, cex.axis=1.25)
axis(4, at = c(300, 350, 400), labels = c(400, 450, 500), las=1, cex.axis=1.25)
abline(h=225, lty=2, col="gray")
bplot(m1.meds[,c(18:24,2)], use.cols = TRUE, outline = FALSE, staplewex=0, col= wes_palette("Royal1")[2],
      boxwex=.15, lty=1, pos = seq(from=.83, by=2, length=8), las=1, xaxt="n", yaxt="n", add=TRUE)
bplot(m2.meds[,c(18:24,2)], use.cols = TRUE, outline = FALSE, staplewex=0, col= wes_palette("Royal1")[3],
      boxwex=.15, lty=1, pos = seq(from=1.17, by=2, length=8), las=1, xaxt="n", yaxt="n", add=TRUE)
bplot(m3.meds[,c(18:24,2)], use.cols = TRUE, outline = FALSE, staplewex=0, col= wes_palette("Royal1")[4],
      boxwex=.15, lty=1, pos = seq(from=1.5, by=2, length=8), las=1, xaxt="n", yaxt="n", add=TRUE)
bplot(data.frame(136, 139, 141, 143, 144, 145, 145, 300), xaxt="n", yaxt="n",
      use.cols = TRUE, boxwex=.8, border= "red", las=1, pos = seq(from=1, by=2, length=8),
      add=TRUE)
axis(1, at = c(1,3,5,7,9,11,13,15), labels = c(expression(N[1]), expression(N[2]), expression(N[3]), expression(N[4]), expression(N[5]), expression(N[6]), expression(N[7]), "Nsuper"), cex.axis=1)
legend("top", legend = c("d", "m1", "m2", "m3" ), fill = wes_palette("Royal1"),
       bty="n")
title(main = "a", line=-1, adj=0.01, cex.main=1.5)

bplot(d.meds[,25:30], use.cols = TRUE, outline = FALSE, staplewex=0, col= wes_palette("Royal1")[1],
      boxwex=.15, lty=1, pos = seq(from=.5, by=2, length=6), xaxt="n",
      ylab = expression(paste("Population change (", lambda, ")"), sep=""),
      xlim=c(0,12), ylim = c(0.2, 1.7), las=1, cex.lab=1.25, cex.axis=1)
bplot(m1.meds[,25:30], use.cols = TRUE, outline = FALSE, staplewex=0, col= wes_palette("Royal1")[2],
      boxwex=.15, lty=1, pos = seq(from=.83, by=2, length=6), las=1, xaxt="n", add=TRUE)
bplot(m2.meds[,25:30], use.cols = TRUE, outline = FALSE, staplewex=0, col= wes_palette("Royal1")[3],
      boxwex=.15, lty=1, pos = seq(from=1.17, by=2, length=6), las=1, xaxt="n", add=TRUE)
bplot(m3.meds[,25:30], use.cols = TRUE, outline = FALSE, staplewex=0, col= wes_palette("Royal1")[4],
      boxwex=.15, lty=1, pos = seq(from=1.5, by=2, length=6), las=1, xaxt="n", add=TRUE)
bplot(data.frame(E_lam[1], E_lam[2], E_lam[3], E_lam[4], E_lam[5], E_lam[6]), xaxt="n",
      use.cols = TRUE, boxwex=.8, border= "red", las=1, pos = seq(from=1, by=2, length=6),
      add=TRUE)
axis(1, at = c(1,3,5,7,9,11), labels = c(expression(italic(lambda[1])),
                                                  expression(italic(lambda[2])),
                                                  expression(italic(lambda[3])),
                                                  expression(italic(lambda[4])),
                                                  expression(italic(lambda[5])),
                                                  expression(italic(lambda[6]))), cex.axis=1)
title(main="b", line=-1, adj=0.01, cex.main=1.5)


bplot(d.meds[,12:17], use.cols = TRUE, outline = FALSE, staplewex=0, col= wes_palette("Royal1")[1],
      boxwex=.15, lty=1, pos = seq(from=.5, by=2, length=6), xaxt="n",
      ylab = "Recruitment (f)",
      xlim=c(0,12), ylim = c(0, 1), las=1, cex.lab=1.25, cex.axis=1)
bplot(m1.meds[,12:17], use.cols = TRUE, outline = FALSE, staplewex=0, col= wes_palette("Royal1")[2],
      boxwex=.15, lty=1, pos = seq(from=.83, by=2, length=6), las=1, xaxt="n", add=TRUE)
bplot(m2.meds[,12:17], use.cols = TRUE, outline = FALSE, staplewex=0, col= wes_palette("Royal1")[3],
      boxwex=.15, lty=1, pos = seq(from=1.17, by=2, length=6), las=1, xaxt="n", add=TRUE)
bplot(m3.meds[,12:17], use.cols = TRUE, outline = FALSE, staplewex=0, col= wes_palette("Royal1")[4],
      boxwex=.15, lty=1, pos = seq(from=1.5, by=2, length=6), las=1, xaxt="n", add=TRUE)
bplot(data.frame(E_f[1], E_f[2], E_f[3], E_f[4], E_f[5], E_f[6]), xaxt="n",
      use.cols = TRUE, boxwex=.8, border= "red", las=1, pos = seq(from=1, by=2, length=6),
      add=TRUE)
axis(1, at = c(1,3,5,7,9,11), labels = c(expression(italic(f[1])),
                                         expression(italic(f[2])),
                                         expression(italic(f[3])),
                                         expression(italic(f[4])),
                                         expression(italic(f[5])),
                                         expression(italic(f[6]))), cex.axis=1)
title(main="c", line=-1, adj=0.01, cex.main=1.5)
#----------------------------------------------------------------------------
# Create Figure S2

js.out.marg[[5]]$mps100 <- js.out.marg[[5]]$rt/js.out.marg[[5]]$min.ess*100
js.out.marg[[5]]$mps1000 <- js.out.marg[[5]]$rt/js.out.marg[[5]]$min.ess*1000

par(family = "serif", mfrow=c(1,1), mar = c(4,5,0,0)) # family = "CM Roman",
boxplot(js.out.marg[[5]]$mps1000 ~ js.out.marg[[5]]$model, log = 'y', xlab= "Model",
        ylab = "Time to 1000 effective samples (min)",
        xaxt= "n", yaxt="n", outline=FALSE, lty=1, staplewex=0)
axis(2, at = c(0, 0.1, 1, 10, 100), las = 1)
axis(1, at = c(1, 2, 3, 4), labels = c("d", "m1", "m2", "m3"), las = 1)
```
